# Profiling Human CMV-specific T cell responses reveals novel immunogenic ORFs

**DOI:** 10.1101/2021.06.10.447997

**Authors:** Rekha Dhanwani, Sandeep Kumar Dhanda, John Pham, Gregory P. Williams, John Sidney, Alba Grifoni, Gaelle Picarda, Cecilia S. Lindestam Arlehamn, Alessandro Sette, Chris A Benedict

**Author notes:** Corresponding authors Alessandro Sette Chris A. Benedict. Present Address: Division of Extramural Activities, National Institute of Allergy and Infectious Diseases, National Institute of Health, Rockville, MD 20852, USA. Present Address: St. Jude Children’s Research Hospital, Arlington, VA 22203.

## Abstract

Despite the prevalence and medical significance of human cytomegalovirus (HCMV) infections, a systematic analysis of the targets of T cell recognition in humans that spans the entire genome and includes recently described potential novel ORFs is not available. Here, we screened a library of epitopes predicted to bind HLA class II that spans over 350 different HCMV ORFs and includes ∼150 previously described and ∼200 recently described potential novel ORFs using an ex vivo IFNγ fluorospot assay. We identified 235 unique HCMV specific epitopes derived from 100 ORFs, some previously described as immunodominant and others that were not previously described to be immunogenic. Of those, 41 belong to the set of recently reported novel ORFs, thus providing evidence that at least some of these are actually expressed *in vivo* in humans. These data reveal that the breadth of the human T cell response to HCMV is much greater than previously thought. The ORFs and epitopes identified will help elucidate how T cell immunity relates to HCMV pathogenesis and instruct ongoing HCMV vaccine research.

**Importance:** To understand the crucial role of adaptive immunity in controlling cytomegalovirus infection and disease, we systematically analyzed the CMV ’ORFeome’ to identify new CMV epitopes targeted primarily by CD4 T cells in humans. Our study identified >200 new T cell epitopes derived from both canonical and novel ORFs, highlighting the substantial breadth of anti-CMV T cell response and providing new targets for vaccine design.

## Introduction

Human cytomegalovirus (HCMV, HHV-5) is a β-herpesvirus that infects the majority of the world’s population. Infection in healthy individuals is characterized by a primary asymptomatic phase followed by the establishment of lifelong persistence/latency in several cell types (1, 2). HCMV’s 236 kbp double stranded DNA genome facilitates its persistence and reactivation when immunity is compromised, with both viral and cellular proteins controlling viral gene expression and regulating the dynamic and reversible latent-lytic cycle that develops over a lifelong infection (3, 4). Although largely persistent, its reactivation in immunocompromised populations, such as transplant recipients and AIDS patients, causes severe disease outcomes (5–11). Congenital infection in the developing fetus is also the leading infectious cause of birth defects (12–18). Moreover, the available antiviral drug therapies are insufficient and often toxic in young children (19–22). Consequently, HCMV is recognized as a major public health problem and development of a vaccine that prevents or at least mitigates virus-induced disease is a top priority (23–25) .

Although both humoral and cell mediated immune responses protect against HCMV infection, a considerable effort has been made towards identifying HCMV targets of CTL responses due to their pivotal role in controlling HCMV disease in immunocompromised individuals (26–29). However, HCMV targets of CD4+ T helper cells, which amplify CTL and antibody responses or may mediate direct antiviral activity themselves, remain to be explored in detail. In order to develop a successful HCMV vaccine, it is imperative to assess the large number of candidate viral proteins for their potential to induce robust CD4+ T cell responses.

Previous work from Sylwester et al. extensively characterized the canonical HCMV proteins that are targeted by CD4+ and CD8+ T cell responses (30), and work by many other groups have identified immunodominant epitopes derived from these that include the 65kDA phosphoprotein (UL83/pp65), immediate early protein 1 (UL123), tegument protein pp150 (UL32), envelope glycoprotein B (UL55), viral transcription factor IE2 (UL122), and major capsid protein (UL86) (31–38). However, a comprehensive analysis of HCMV epitope-specific T cell responses has been challenging, mainly due to the large size of virus and the evolving impact that persistent infection has on the memory pool. Stern-Ginossar et al. recently reported all the HCMV RNAs found to be associated with ribosomes in infected fibroblasts, increasing the potential number of ORFs the virus may encode by ∼3 fold (39). Here, we designed a comprehensive screening approach to assess potential T cell responses against 563 of these ORFs, which included both previously reported and potentially novel HCMV proteins. 2593 15-mer peptides were predicted using computational algorithms, and a high throughput screen was performed using an IFNγ fluorospot assay to identify epitopes targeted by both CD8+ and CD4+ T cells in healthy HCMV-infected adults. This ‘whole ORFeome’ approach resulted in the identification of more than 200 new CD4+ and CD8+ T cell epitopes.

## Results

### Targets of HCMV T cell reactivity

To define the epitopes targeted by HCMV-specific T cell responses in healthy adults, we screened PBMCs of 19 subjects, 10 males and 9 females, recruited from the San Diego blood bank (SDBB). The HCMV seropositivity of all the subjects was confirmed by IgG ELISA **(Fig. S1A)**. We tested a total of 2593 15-mer HCMV peptides covering a total of 563 ORFs (39). Removing the predicted ORFs that were located entirely within longer ORFs resulted in a set of 359 completely unique ORFs. This set consists of approximately 150 “canonical” ORFs, with an additional 200 identified by ribosomal RNA profiling (39). These 15-mer peptides corresponded to epitopes likely to be dominant based on a bioinformatic method that predicts promiscuous binding to HLA class II molecules (40). Each ORF analyzed contained a minimum of 2 predicted epitopes, with the exception of very small ORFs of less than 15-20 amino acid residues, in which case at least one peptide was synthetized. The 2593 peptides were arranged in 89 pools of 28 to 30 15-mers. The PBMC reactivity of each of the 89 pools was assayed directly *ex vivo* using an IFN-γ Fluorospot assay. After identifying the pools that resulted in IFN-γ production in HCMV+ individuals, the top 10 most reactive pools (that, on average, accounted for more than 90% of the reactivity observed within each subject) were then deconvoluted to identify the specific epitopes (**Fig. S2)**. Representative results from the initial screening and the deconvolution of a pool in a representative subject are shown in **Fig. 1A-B.** In conclusion, the results shown here indicate that human T cell responses to HCMV recognize a wide breadth of different epitope specificities.

**Fig. 1.**
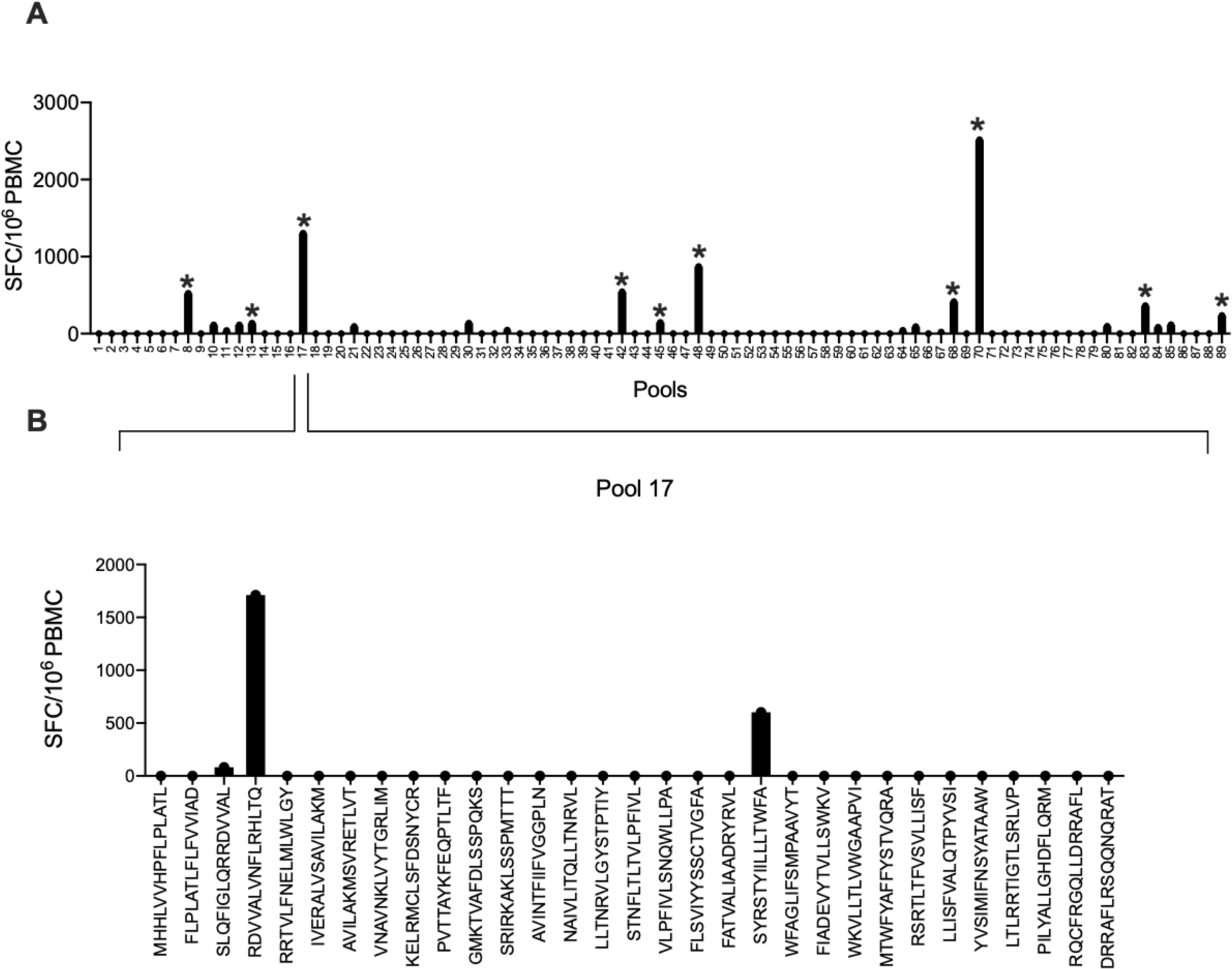
Strategy for HCMV epitope-specific T cell identification: PBMCs from HCMV seropositive subjects were stimulated with 2 µg/ml pools and plated on IFN-γ coated fluorospot plates for 20 hours. The top 10 positive pools (indicated by * on bars) were deconvoluted to identify individual epitopes. PBMC were stimulated with 10 µg/ml of each individual peptide contained in the pool and reactivity was measured by IFN-γ fluorospot assay. (A) SFC/10^6^ PBMC for one representative subject against the 89 peptide pools (B) Deconvoluted pool representing individual peptides.

### Characterization of CMV epitope-specific immune responses

The deconvolution of the top 10 pools from each subject identified widespread reactivity directed against 235 unique epitopes **(Fig. S3 and Table 1)**. Interestingly, females tended to show both a higher frequency and magnitude of epitope-specific responses when compared to males, although this did not reach statistical significance (**Fig. S4)**. On average, each subject recognized 25 epitopes (**Fig. 2A**) and all subjects recognized at least 2 (range 2-57, **Fig. 2B**). Specifically, 6 out of 19 donors recognized 21-30 epitopes. A quarter of the epitopes (58 of the 235 recognized) were recognized by three or more subjects **(Fig. 2C)**, and these accounted for 76% of the total T cell response **(Fig. 2D)**.

**Fig. 2.**
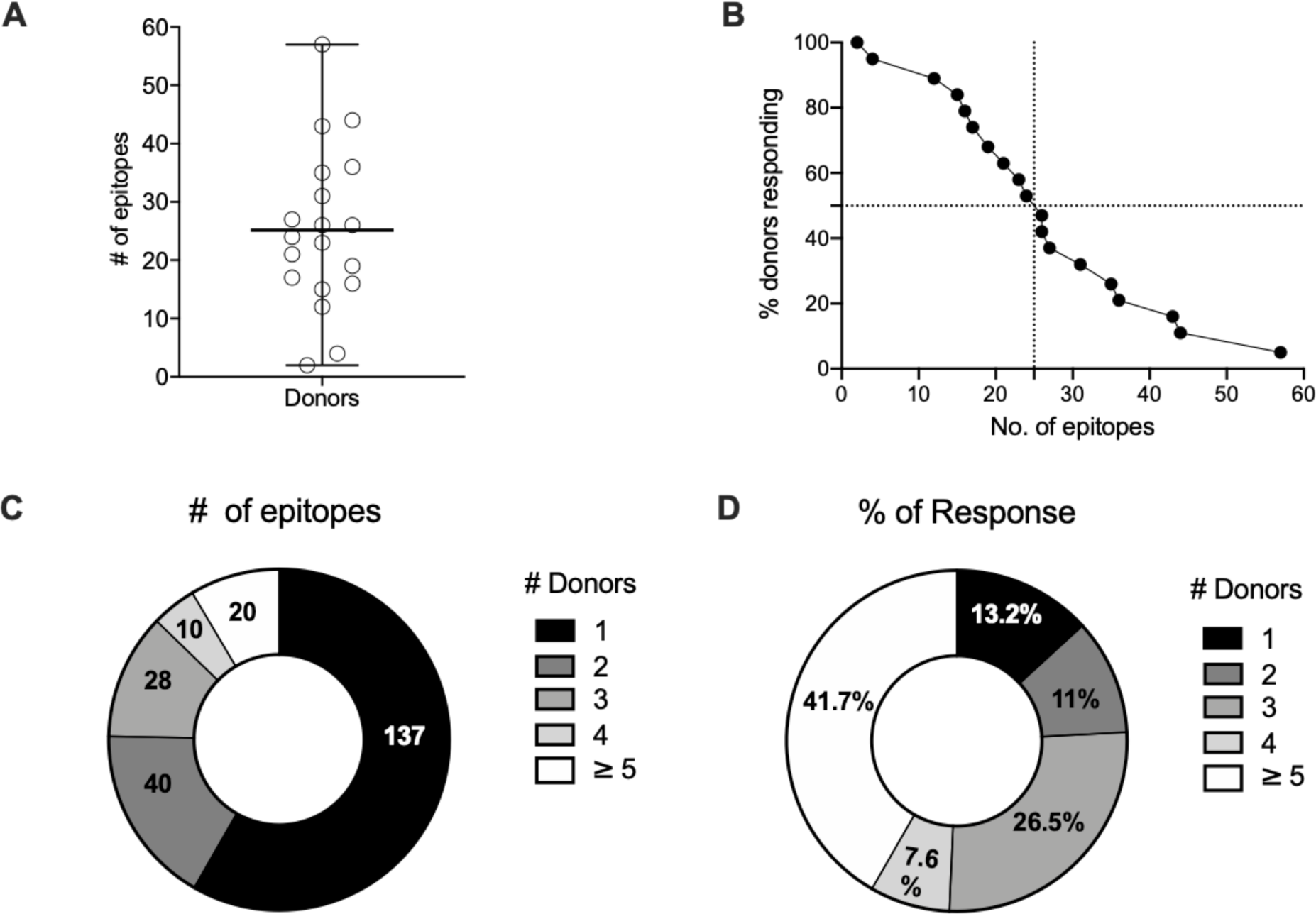
Breadth and dominance of HCMV T cell responses: (A) The number of epitopes recognized by each donor, mean ± range. (B) Proportion of the 19 donors that responded to the indicated number of epitopes. (C) Epitopes by number of responding donors. (D) epitope % of total response by number of responding donors.

**Table 1:**
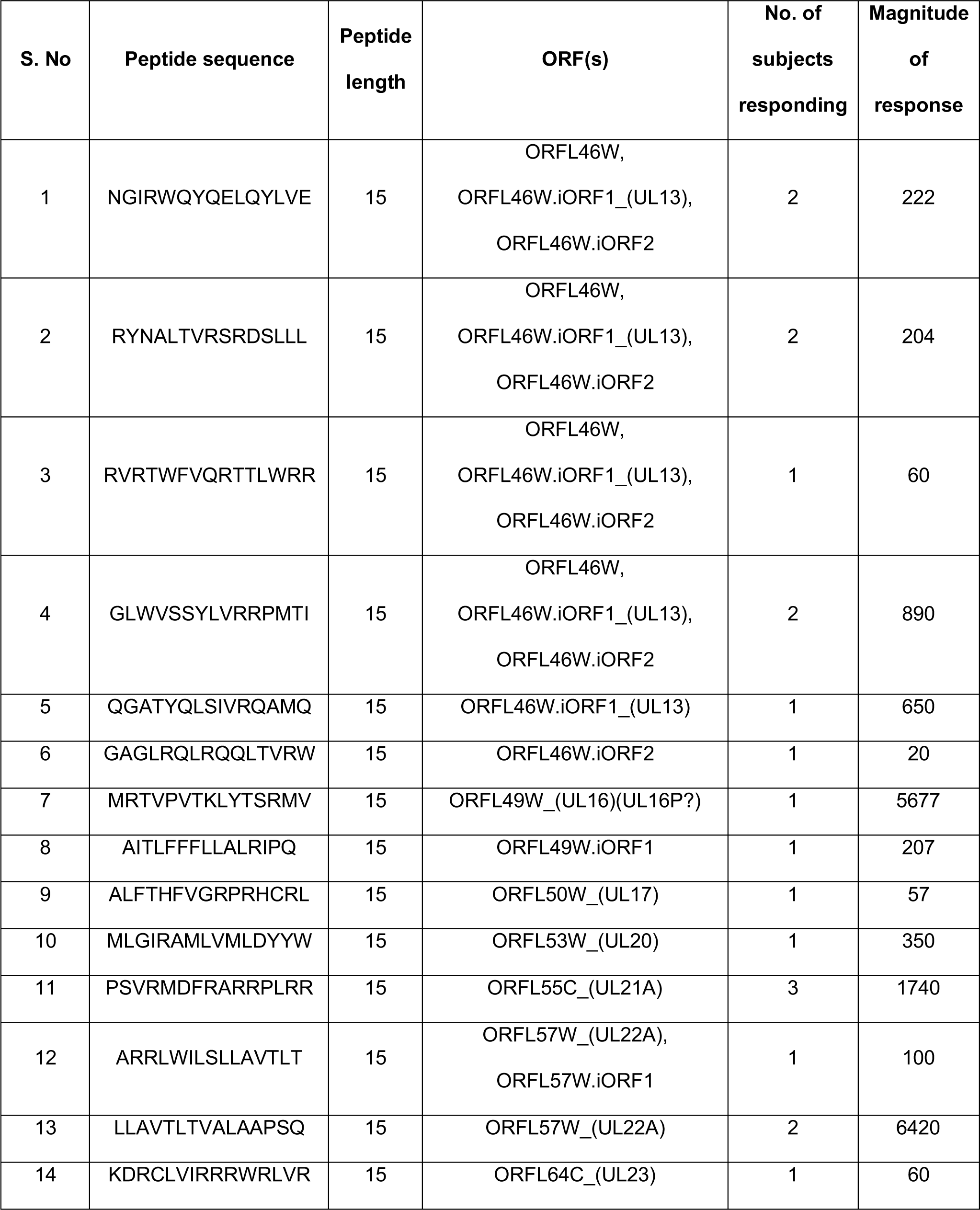
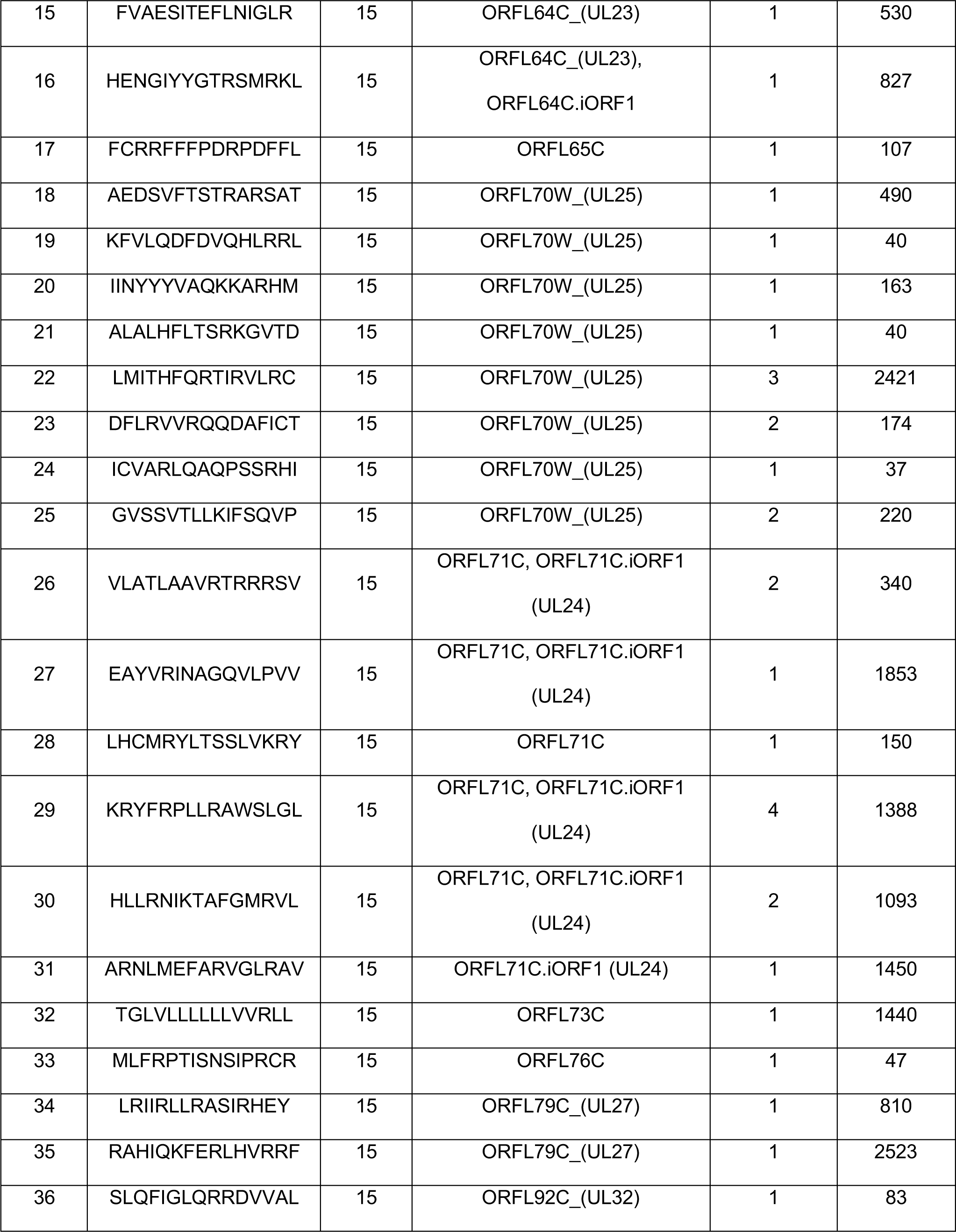
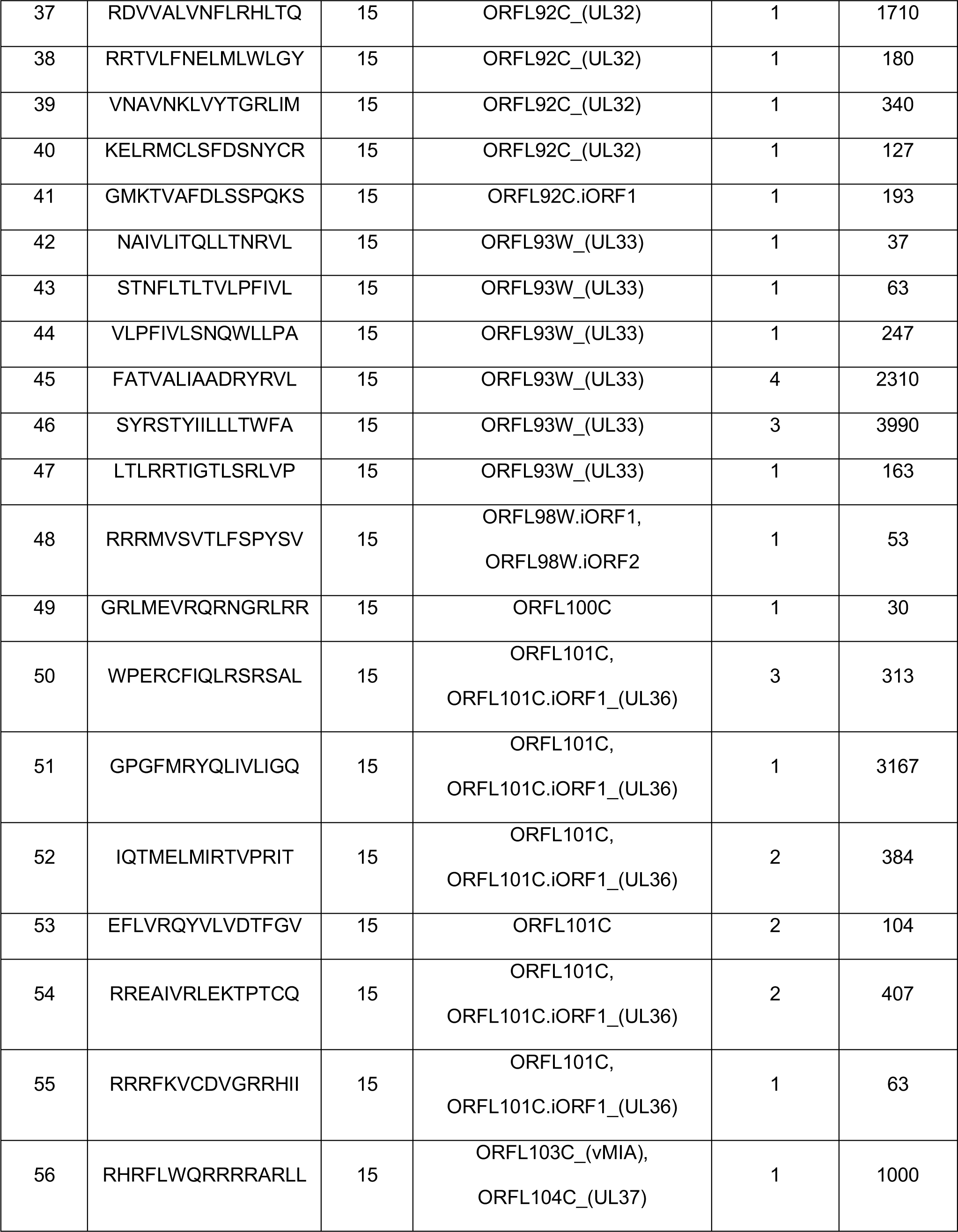
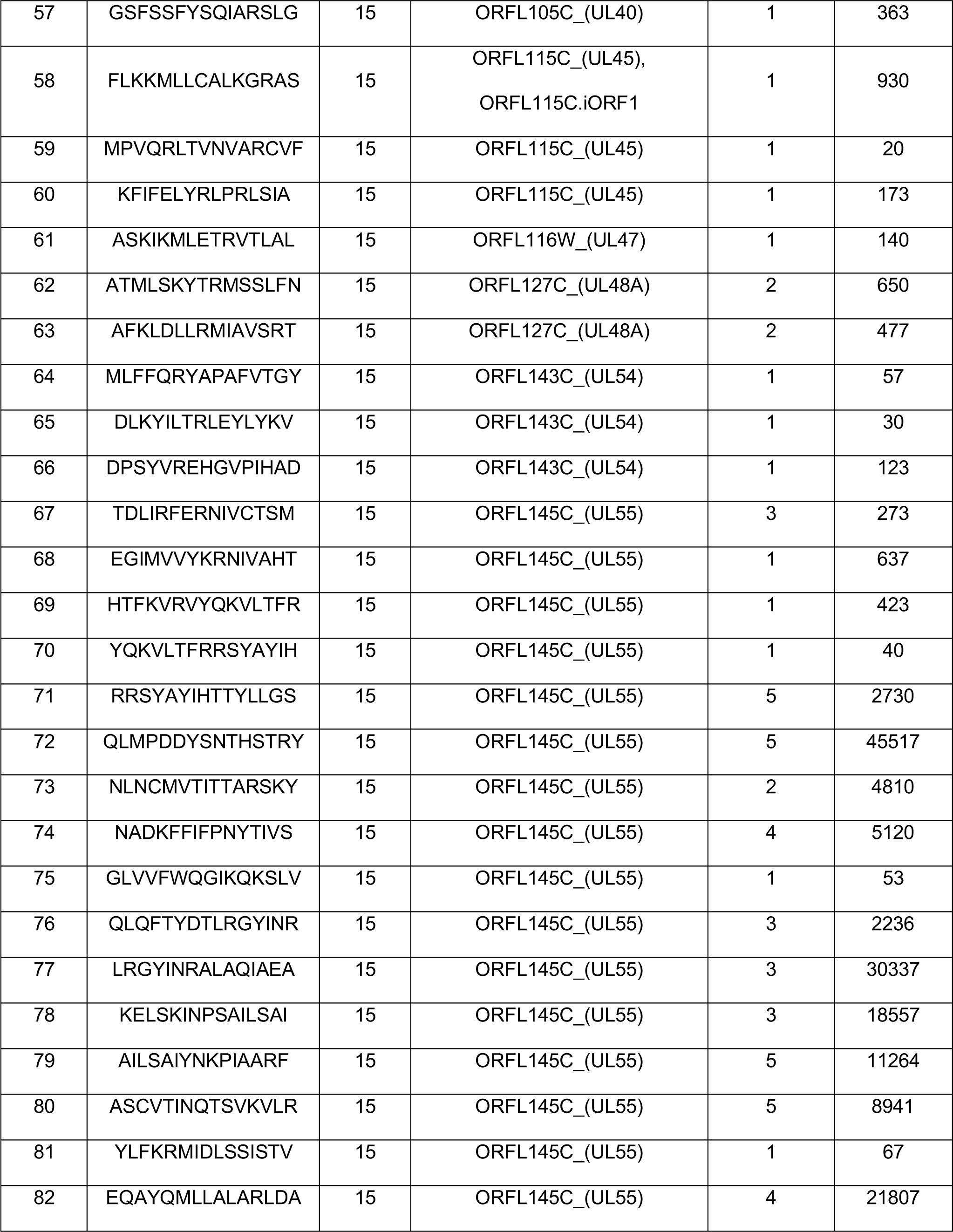
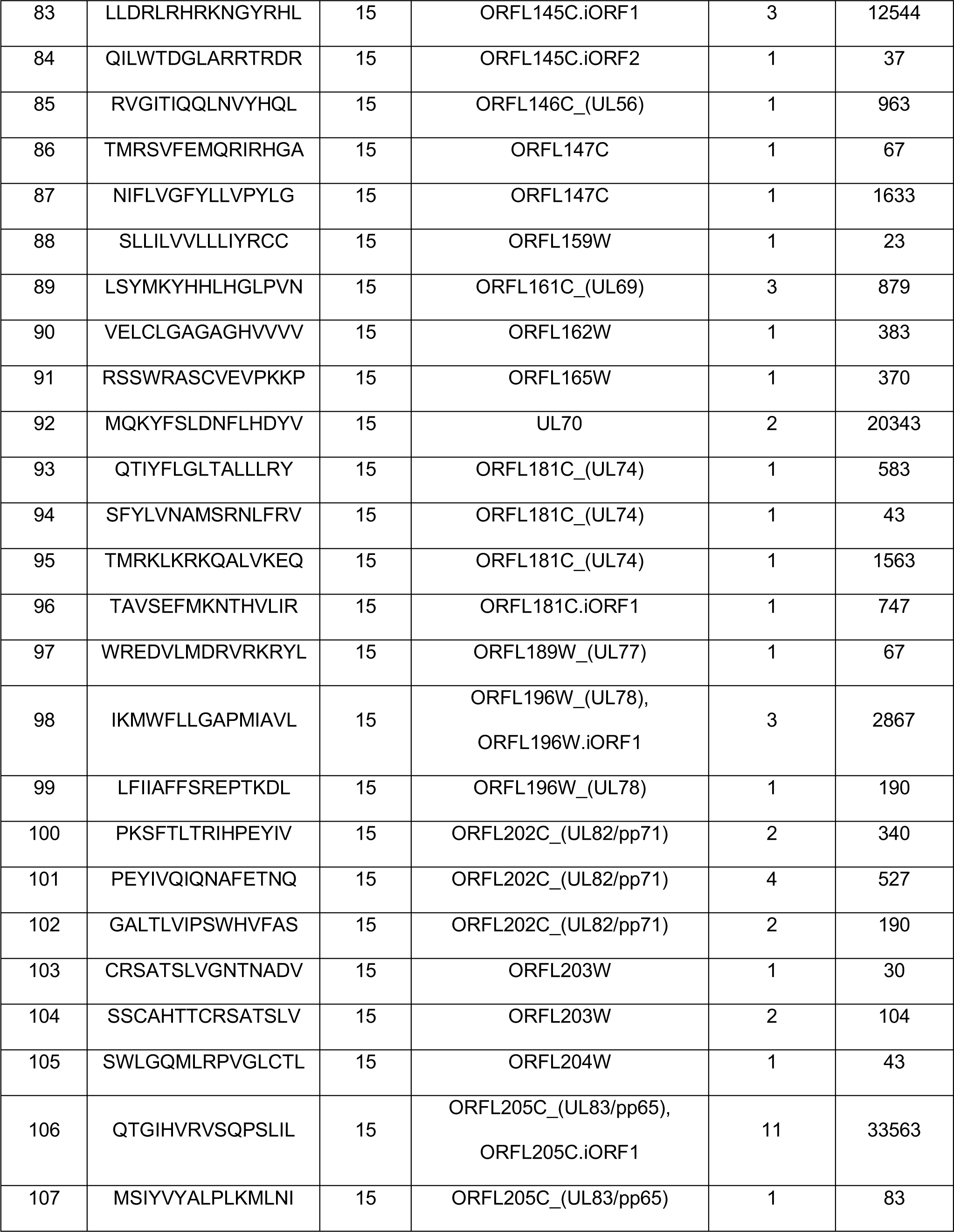
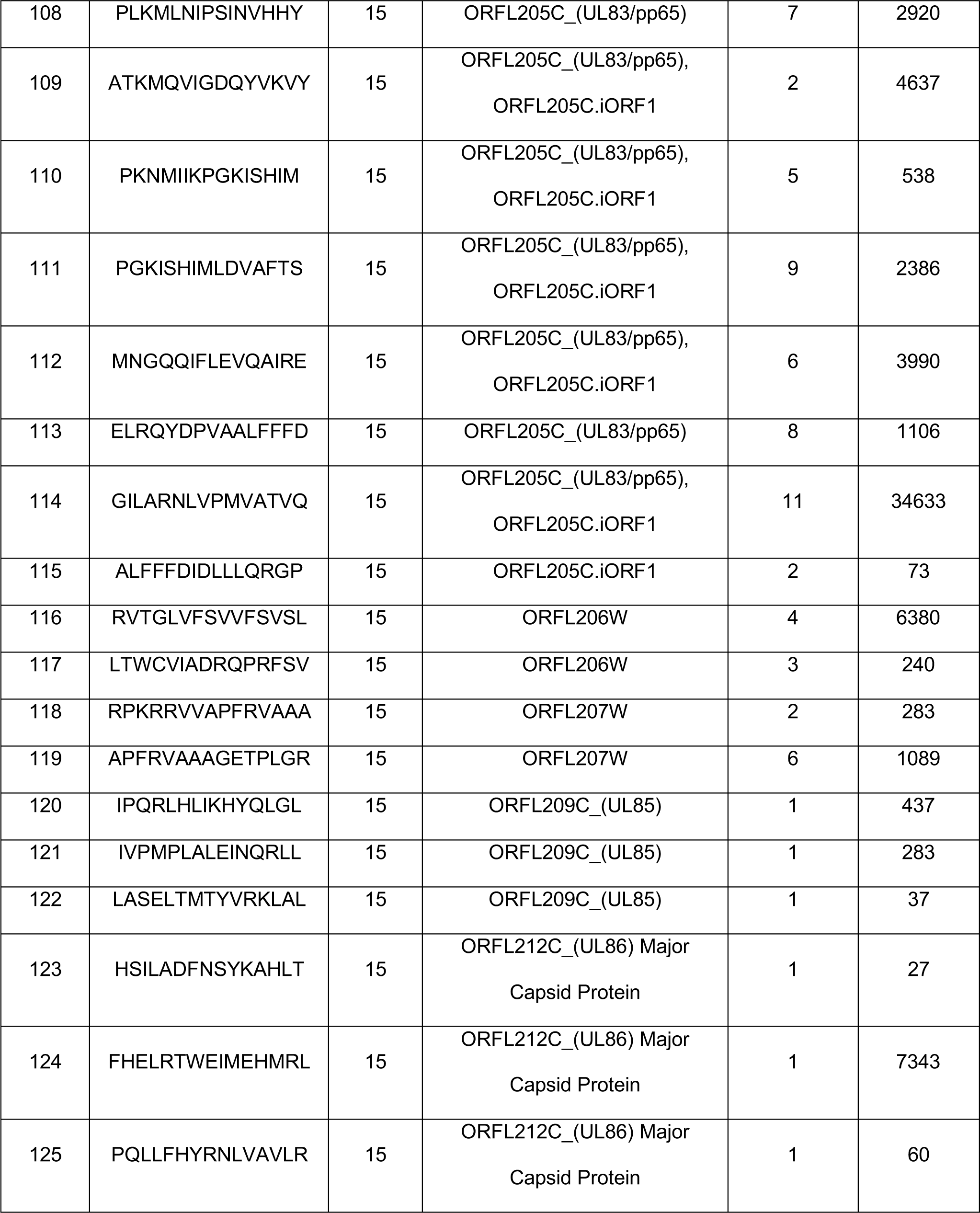
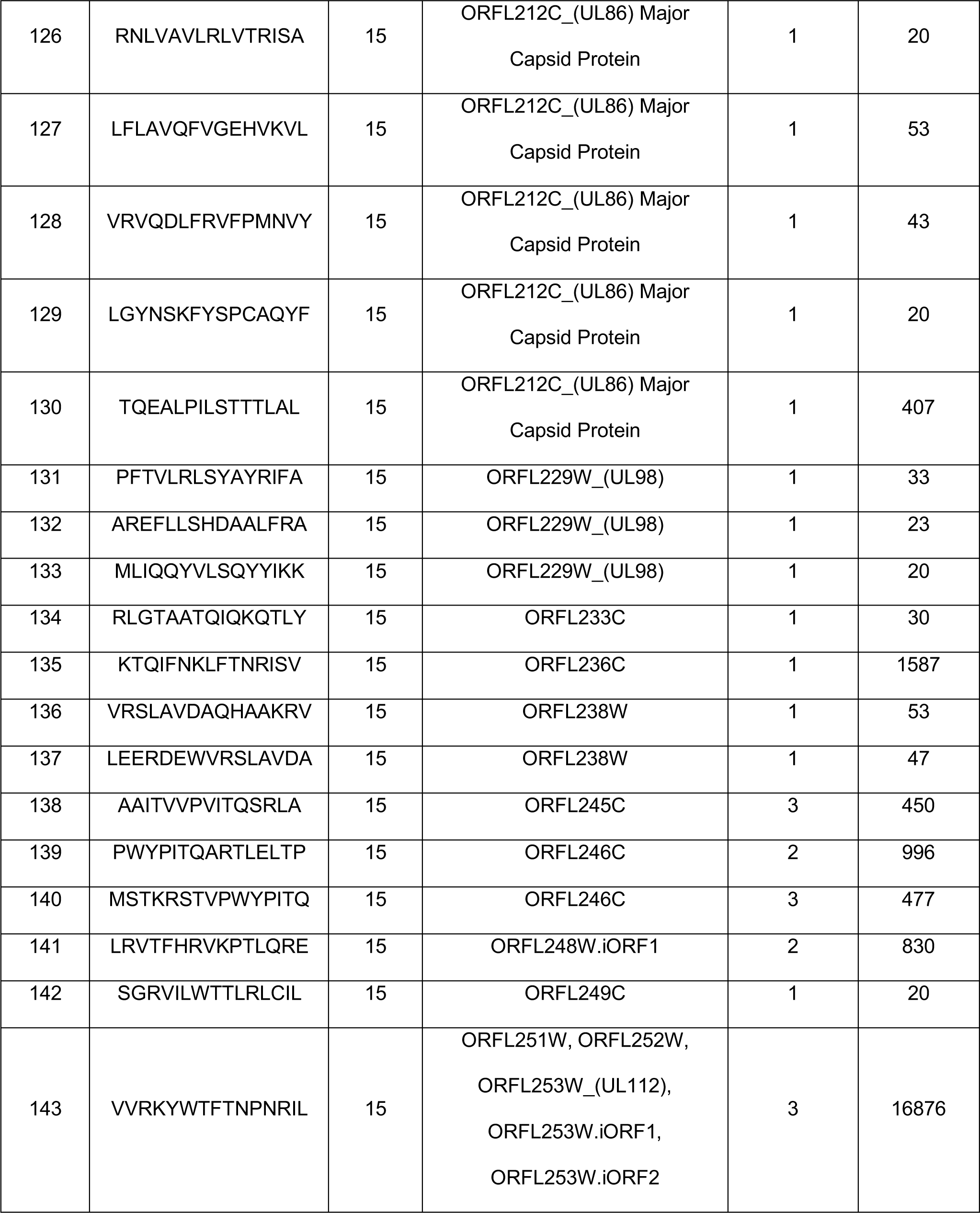
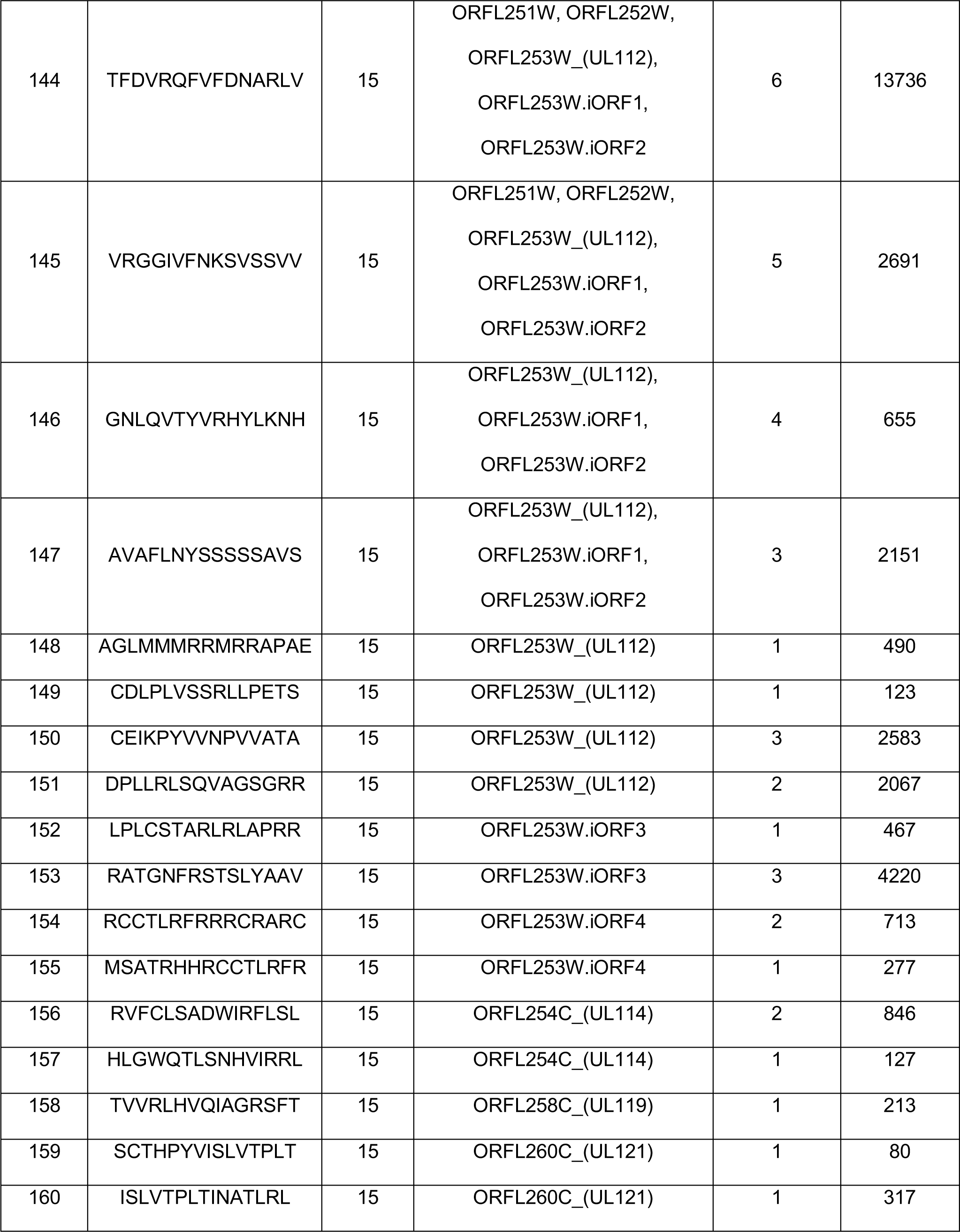
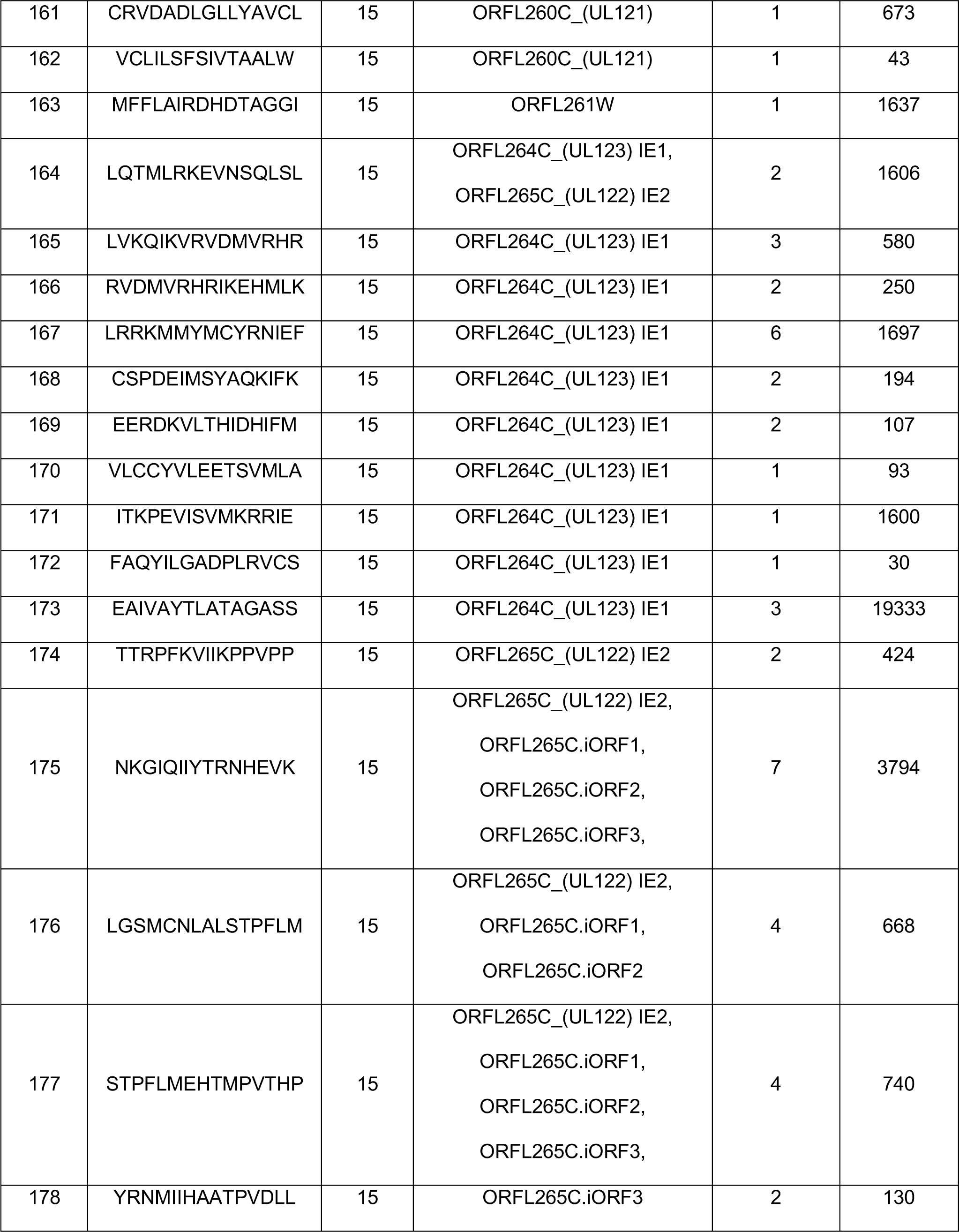
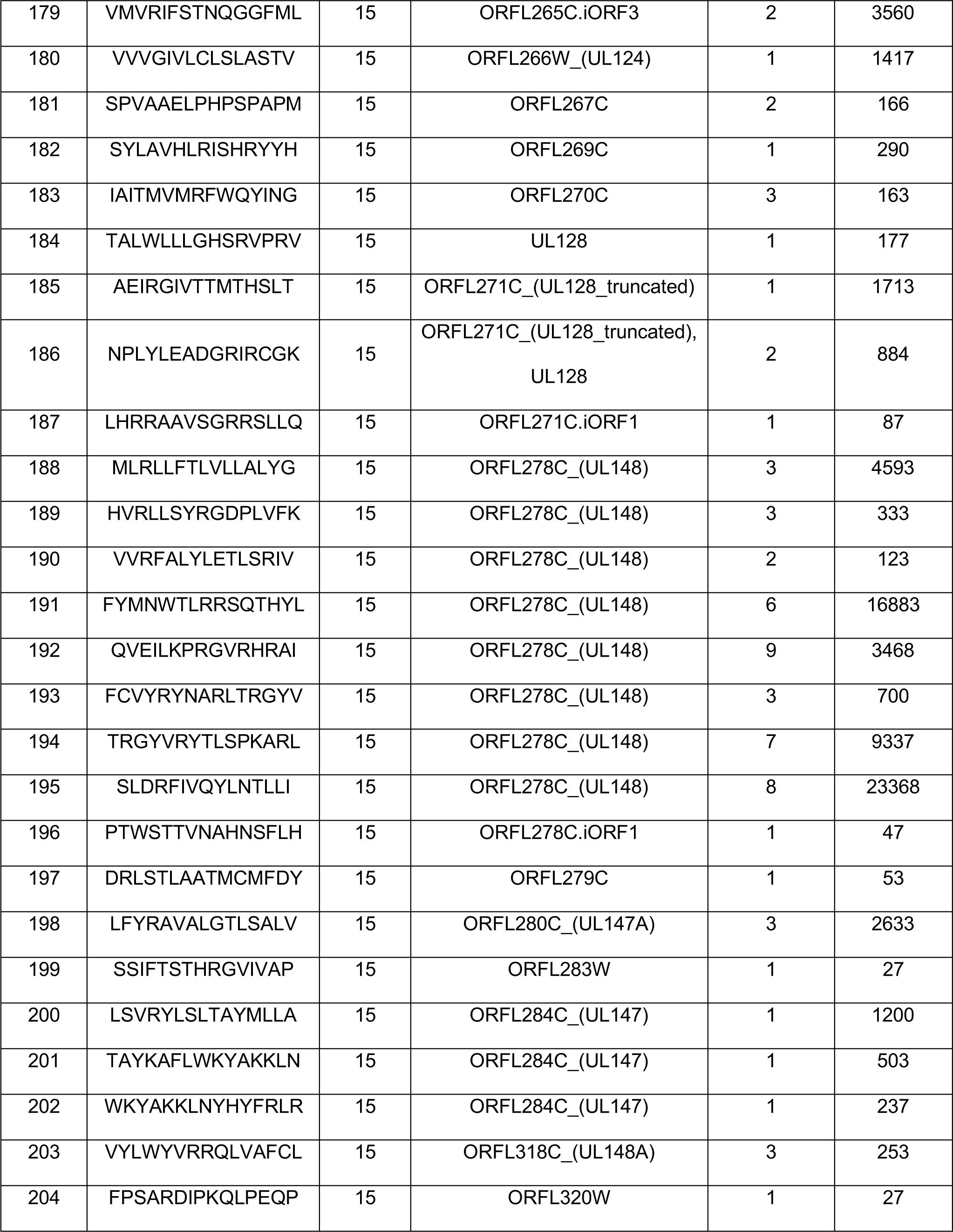
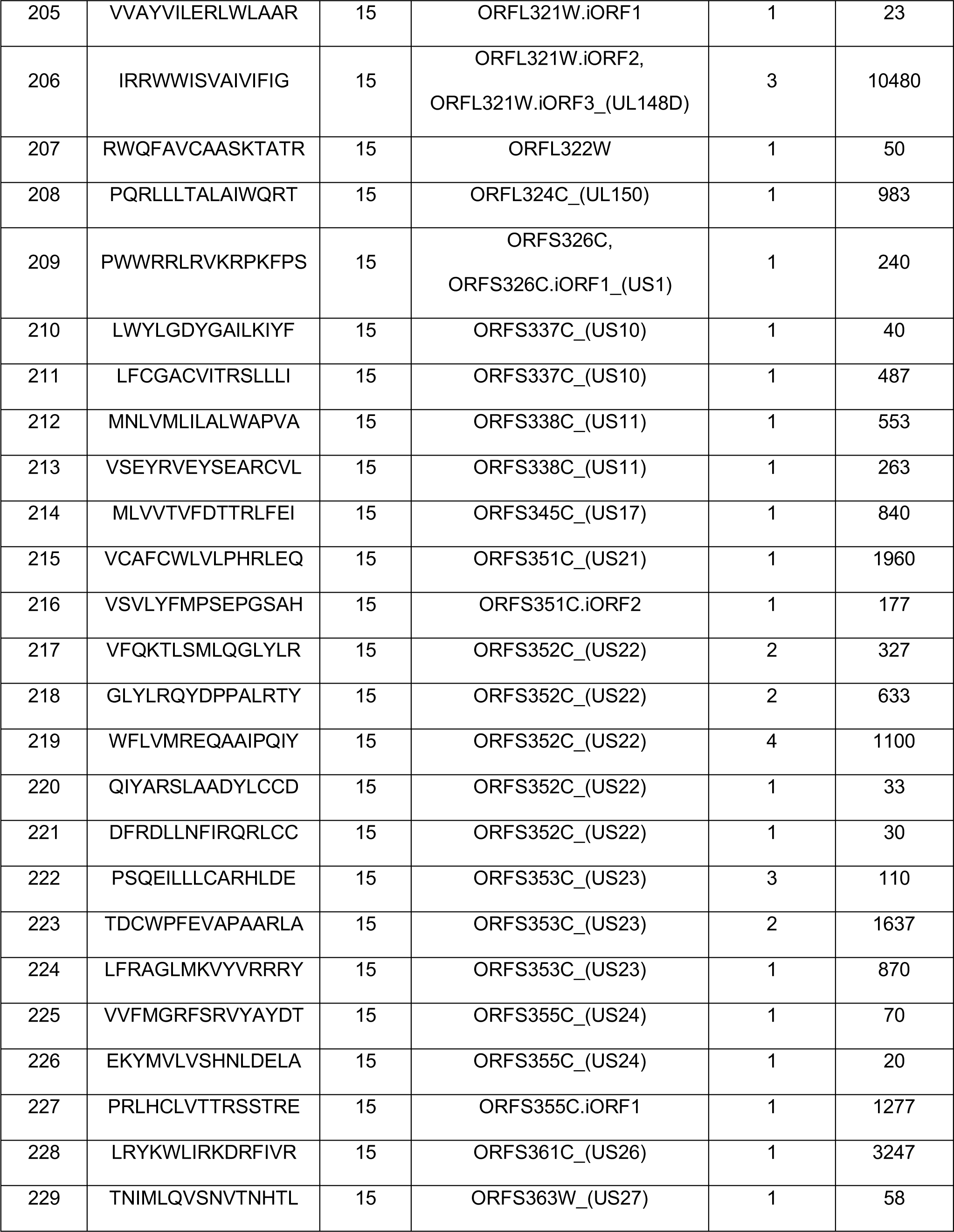
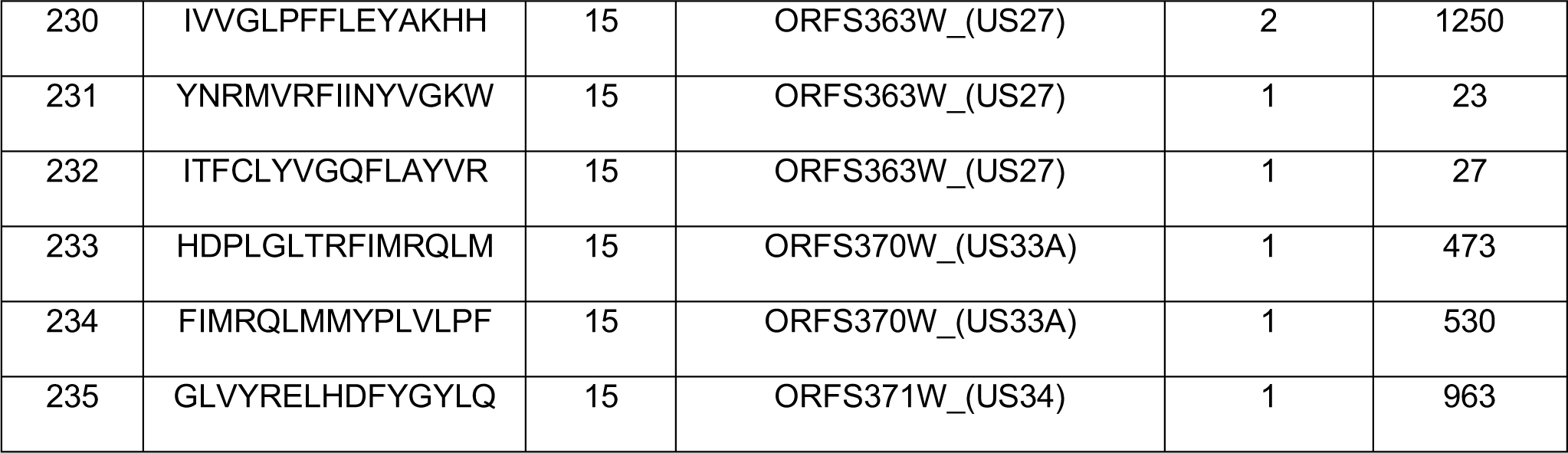
Details of HCMV specific 235 epitopes identified in the screen.

We further characterized the phenotype of the T cell responses directed against these 58 dominant epitopes by intracellular IFN-γ staining (representative results shown in **Fig. 3A**, with the flow cytometry gating strategy shown in **Fig. S5A**). In the majority of tested subjects, the responding T cells were CD4+. More specifically, 68% of all responding T cells were CD4+ and 13% contained both IFN-γ+ CD4+ and CD8+. In 18% of the cases, only CD8+ T cells responded to these 58 epitopes **(Fig. 3B)**. Similarly, if the magnitude of the response was considered, 70% of the IFN-γ response was attributable to CD4+ T cells and only 30% emanated from CD8+ T cells **(Fig. 3C)**. The fact that the responses were dominated by CD4+ T cells is consistent with the fact that the peptides tested were originally selected based on their predicted likelihood to bind HLA class II alleles. In turn, the occasional identification of epitope-specific CD8+ T cell responses in many cases likely reflects class I epitopes nested within the 15-mer epitopes tested in the screen. Overall, these results indicate that, as expected, the screening strategy employed mostly identifies targets of CD4+ T cell reactivity.

**Fig. 3.**
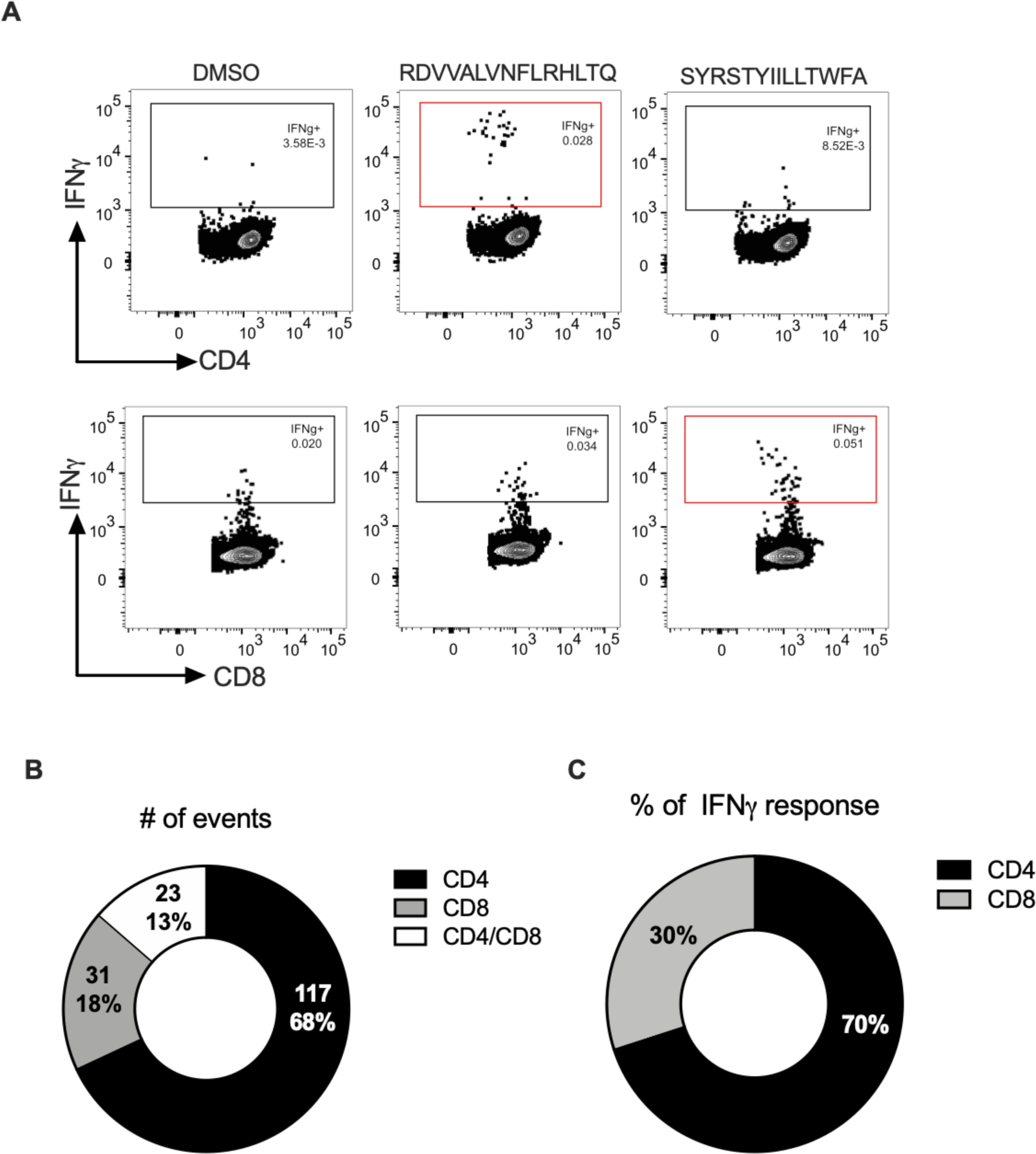
Phenotypic characterization of HCMV T cell responses: (A) Representative FACS plots for intracellular IFN-γ production by CD4+ and CD8+ T cells (gating axis in red) upon stimulation with two of the scoring peptide epitopes that induced them (B and C). The number of events and % response attributable to CD4+ and CD8+ T cell responses of dominant epitopes (n=58) that demonstrated response frequency of 0.15 (15%) (i.e recognized by 3 or more donors).

### Analysis of the ORF of origin of the identified epitopes

The 235 epitopes identified mapped to a total of 100 of the 359 unique ORFs screened. Of those, 28 ORFs contained >3 immunogenic peptides and 18 ORFs were recognized in 15% or more of the donors (**Fig. 4).** Notably, the previously well-characterized immunodominant ORFs such as envelope glycoprotein B (UL55), IE1 (UL123), tegument protein pp65 (UL83), major capsid protein UL86, IE2 (UL122), and pp150 (UL32) were amongst those associated with more than three immunogenic peptides.

**Fig. 4.**
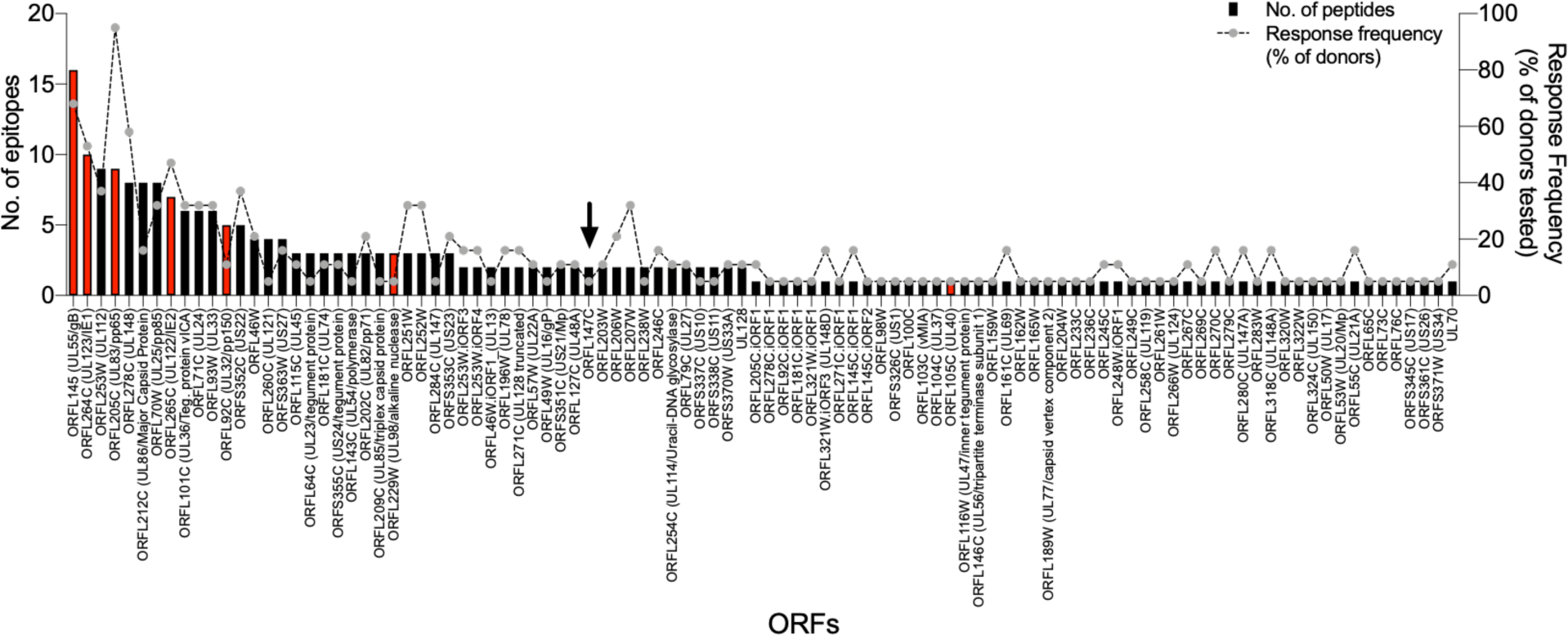
T cell epitope distribution by ORF of origin: 235 epitopes mapped to 100 ORFs. Left Y axis denotes the number of epitopes associated with each ORF (bars) and right Y axis denotes the response frequency associated with each ORF (dotted line). Seven canonical ORFs that were common in IEDB and the present screen are denoted in red. ORFL147C (arrow) is the first ‘novel’ ORF identified by rRNA profiling from left-to-right, and only induces responses in 2/19 individuals tested.

To address the novelty of our findings, we compared our results with ORFs that have already been reported and curated in the Immune Epitope Database (IEDB http://www.iedb.org) (41), as a source of defined epitopes. Specifically, a query of the IEDB in October 2020 for previously characterized targets of T cell responses tested in at least 19 donors and with a minimum response frequency of 15% revealed 7 ORFs that match the conditions of our screening results: UL83/pp65 (ORFL205C), UL123/IE1 (ORFL264C), UL122/IE2 (ORFL265C), UL55/gB (ORFL145C), UL32/pp150 (ORFL92C), UL40 (ORFL105C) and UL98 (ORFL229W) **(Fig. 4)**.

The same query revealed three additional ORFs that were not identified in our screen. These ORFs were associated with a limited number of literature-reported and IEDB curated epitopes: UL75/gH (ORFL184C; 1 epitope), UL44/DNA-pol (ORFL112C.iORF1; 3 epitopes) and UL138 (ORFL313C; 1 epitope). Importantly, our screen identified 93 ORFs that were not previously described as targets of T cell responses **(Fig. 5).**

**Fig. 5.**
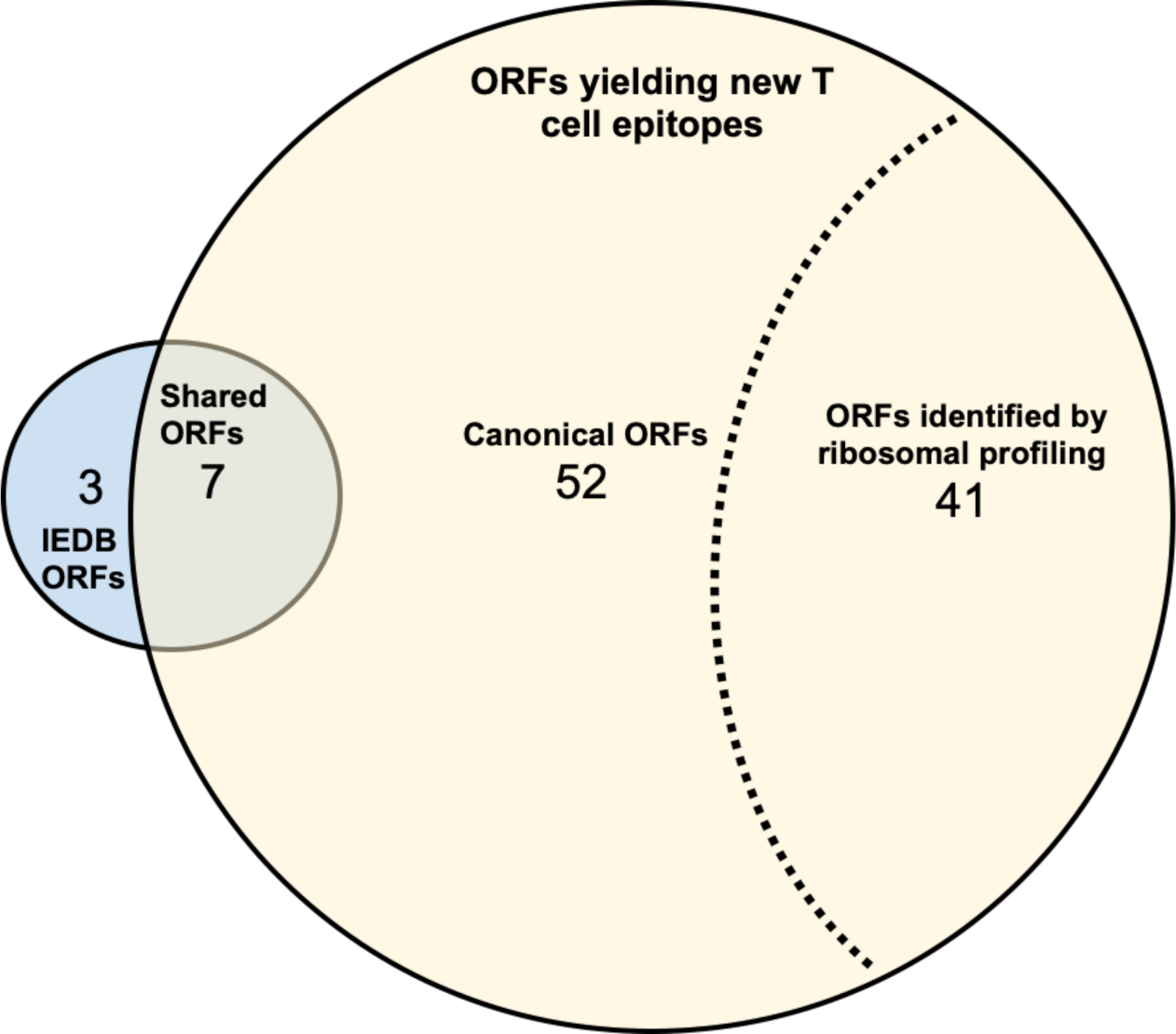
Overlap between IEDB-reported and new immunogenic ORFs identified in this T cell epitope screen. New epitopes were identified in all 100 ORFS, including 7 ORFs previously reported in the IEDB to be targets of T cell responses. Of the 93 ORFs found to be new targets of T cells, 52 were canonical and 41 were ‘novel’ as identified by recent ribosomal mRNA profiling studies.

Notably, 52 of these 93 ORFs were already described in the ‘canonical HCMV’ annotated genome, but not all have been described as targets of human T cell responses. Even more strikingly, 41 of these 93 ORFs corresponded to those viral mRNAs only identified by recent ribosomal profiling studies (39), providing evidence that they are translated in HCMV infected cells. These results indicate that our approach successfully re-identified known ORFs as targets of T cell responses, and perhaps most importantly, greatly expanded the repertoire of canonical and ‘novel’ ORFs recognized by T cells in healthy adults.

### Novel identified epitope pools elicit antigen specific CD4+ T cell responses

Lastly, we wanted to explore whether the epitopes identified in the presented study could, alone or in combination with previously described epitopes, be utilized to generate epitope “MegaPools’” (MP) (42–46) to allow detection of CMV-specific CD4 T cell responses. Accordingly, we generated a ‘P235’ MP encompassing the 235 CMV epitopes identified in the present study. As a comparison, we considered the commercially available CMV peptide pool (Mabtech, catalog 3619-1) encompassing a total of 42 CD4 and CD8 epitopes. Additionally, we synthetized a MP of known class II epitopes curated in the IEDB database, encompassing a total of 187 CD4 epitopes (IEDB-II, Table 2).

**Table 2:**
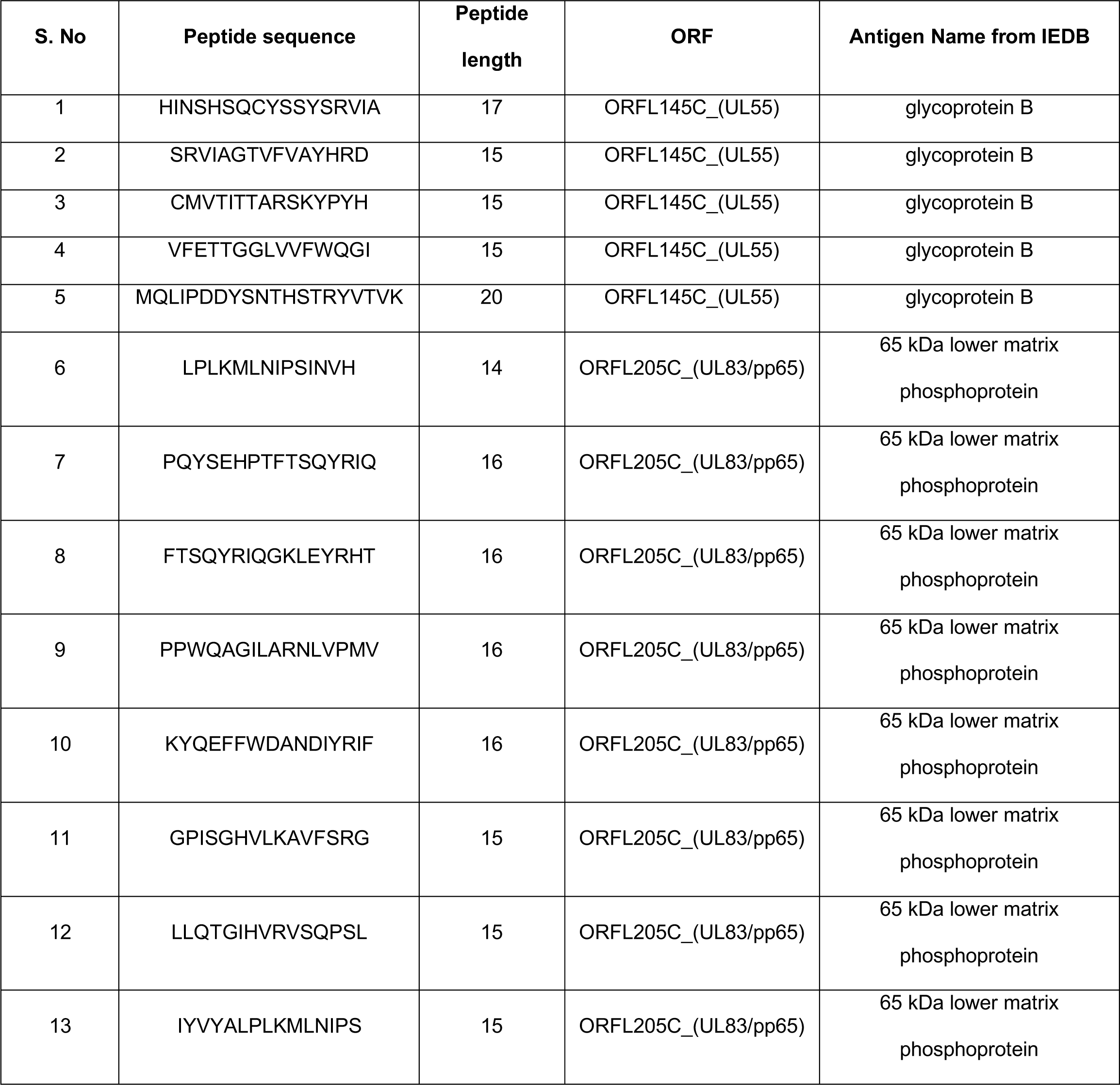
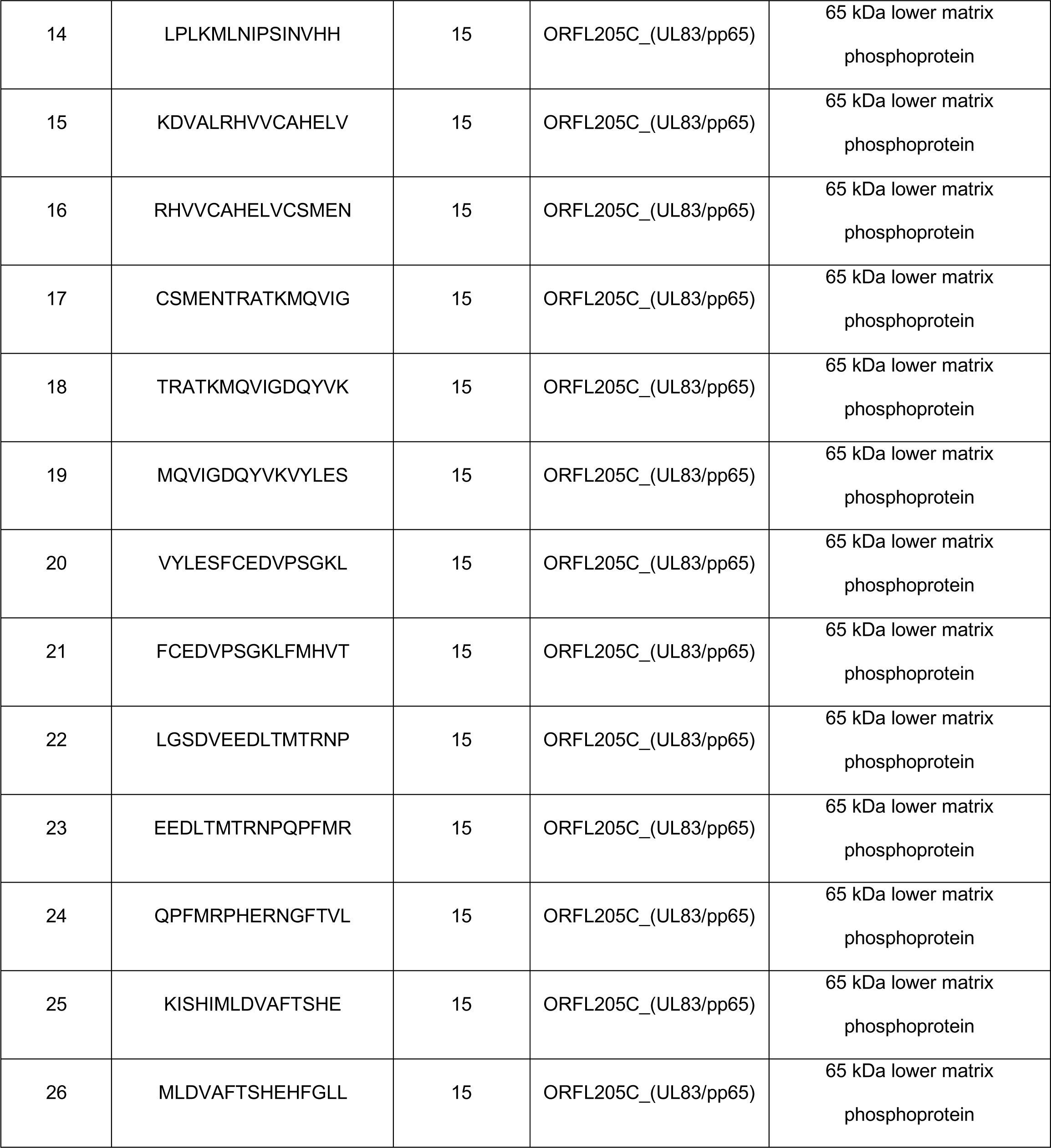
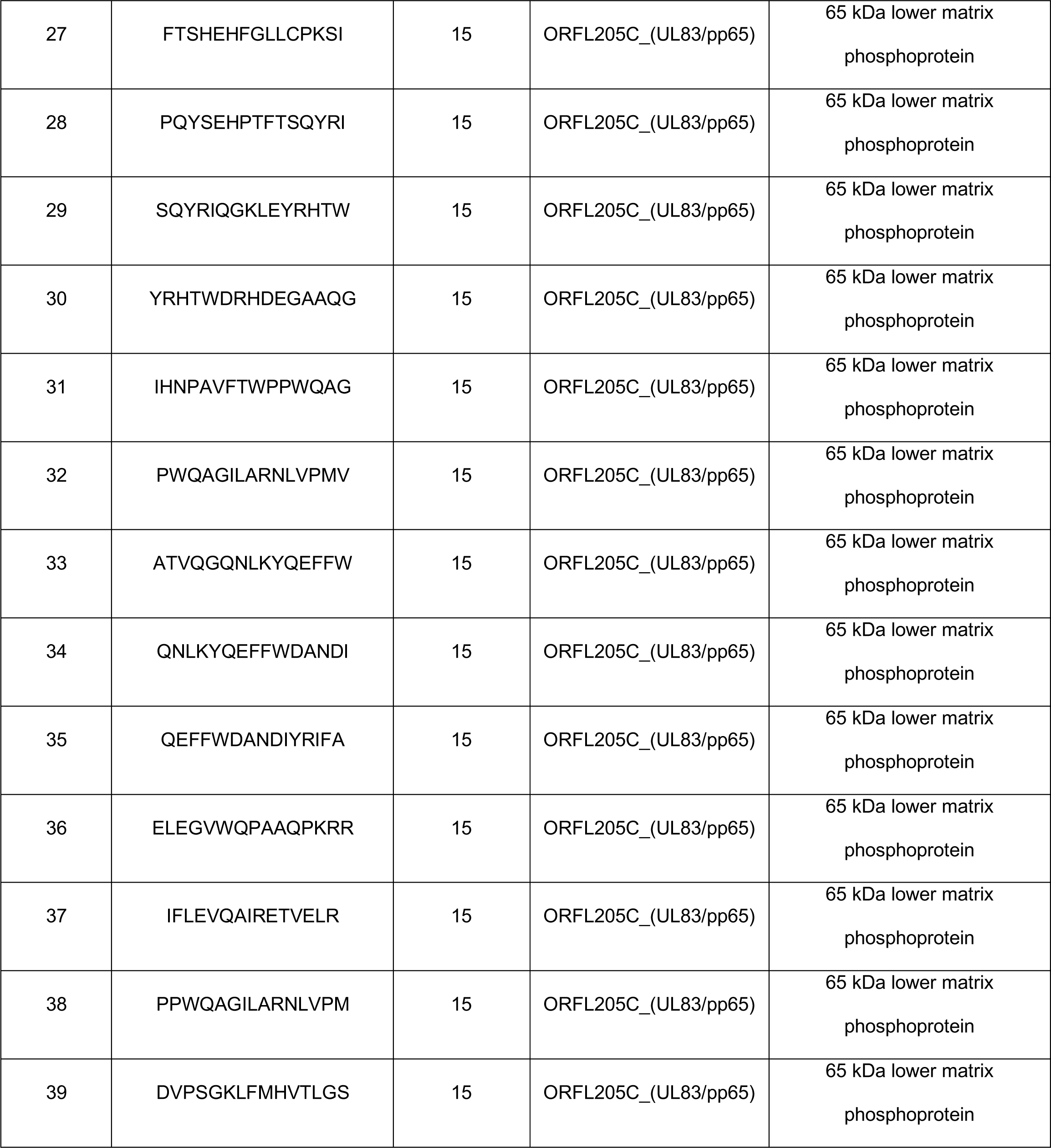
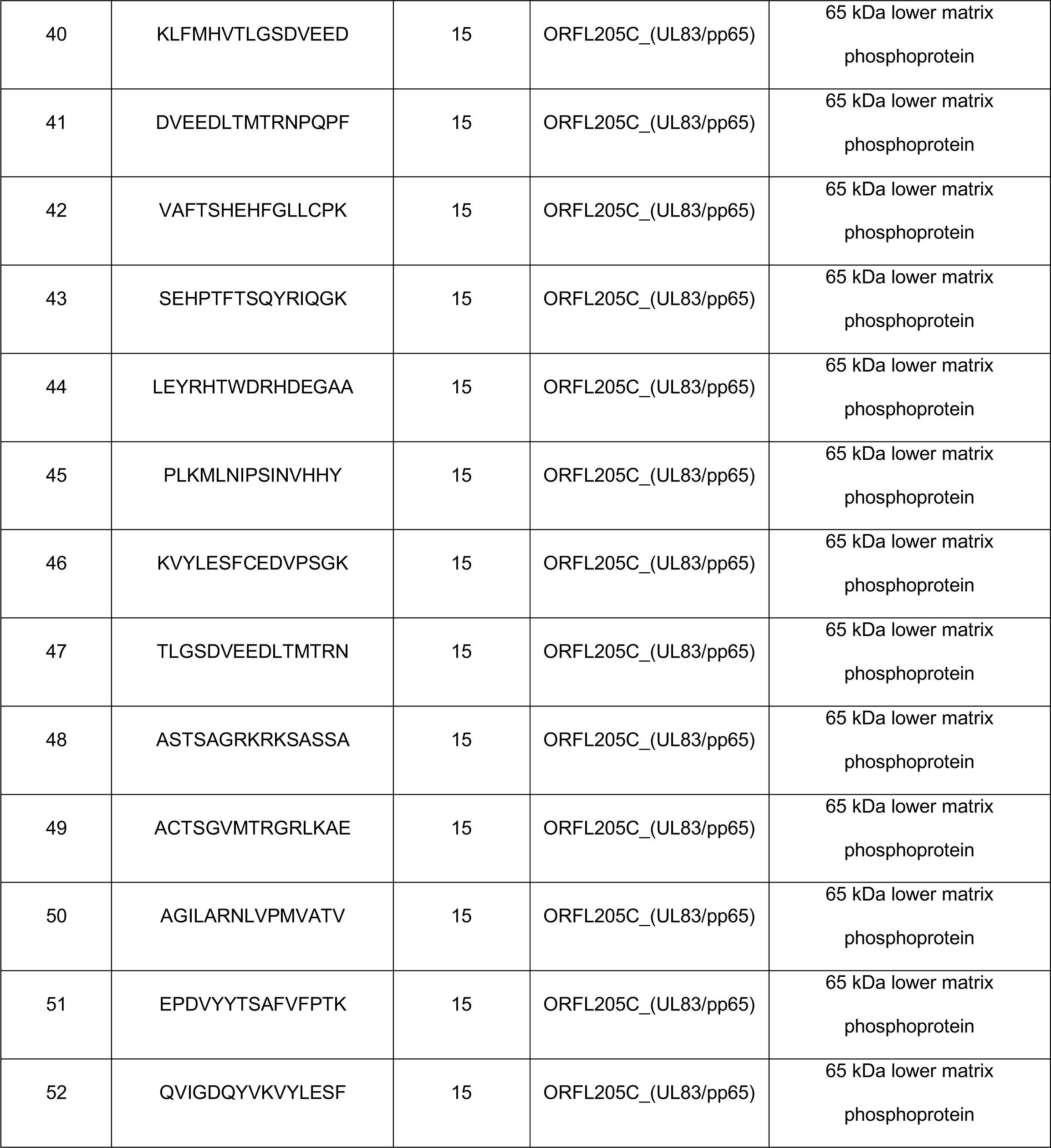
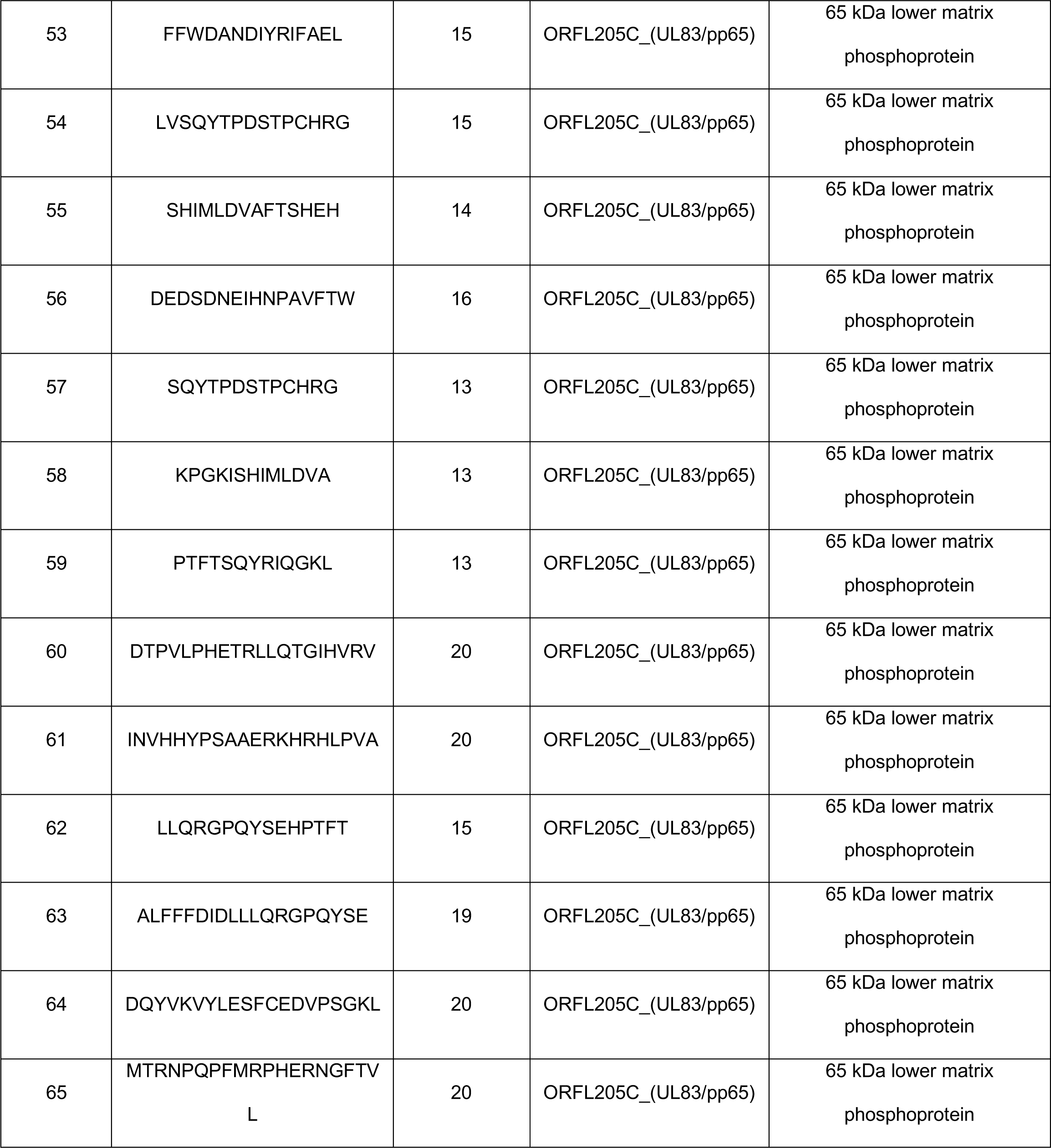
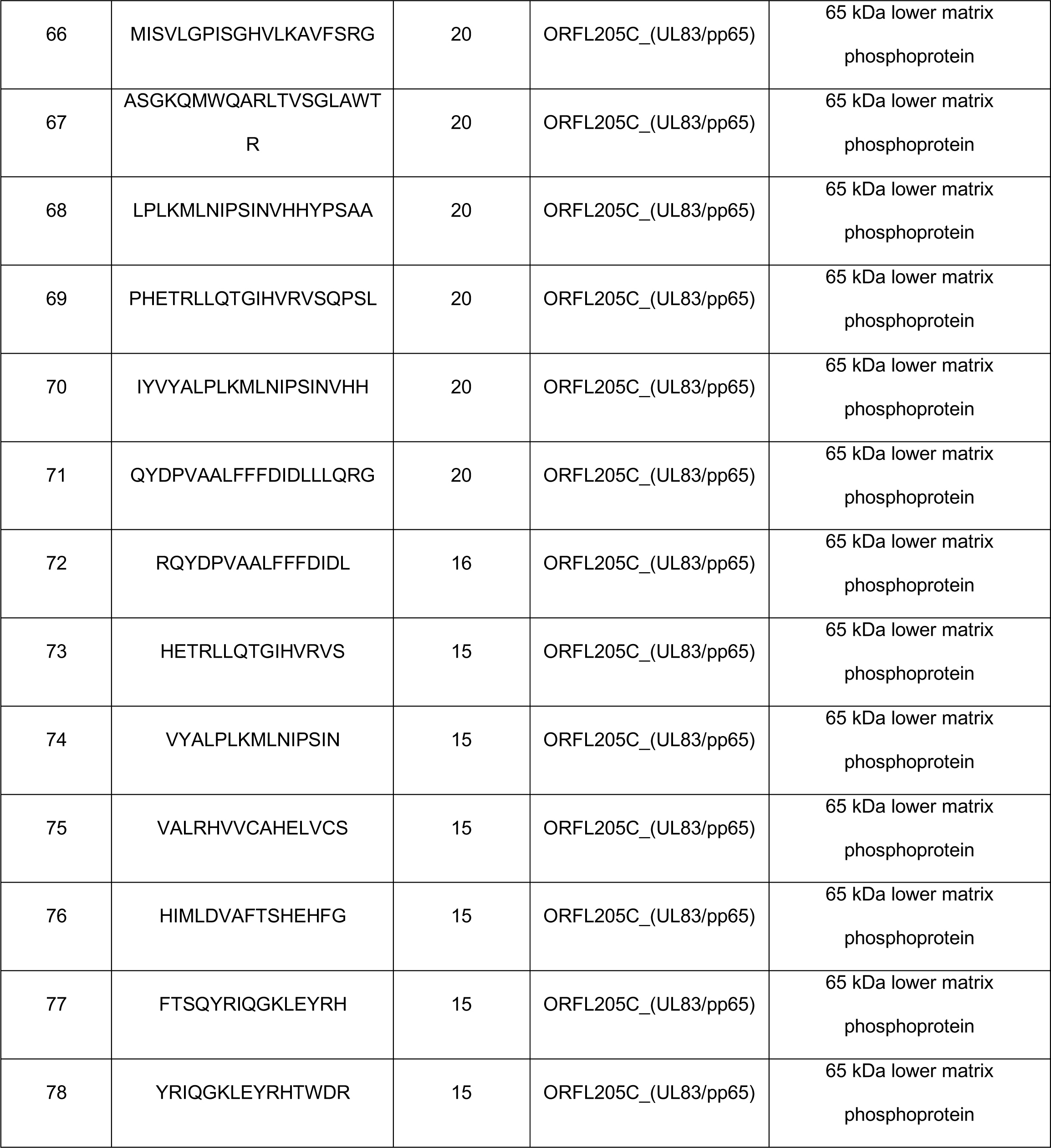
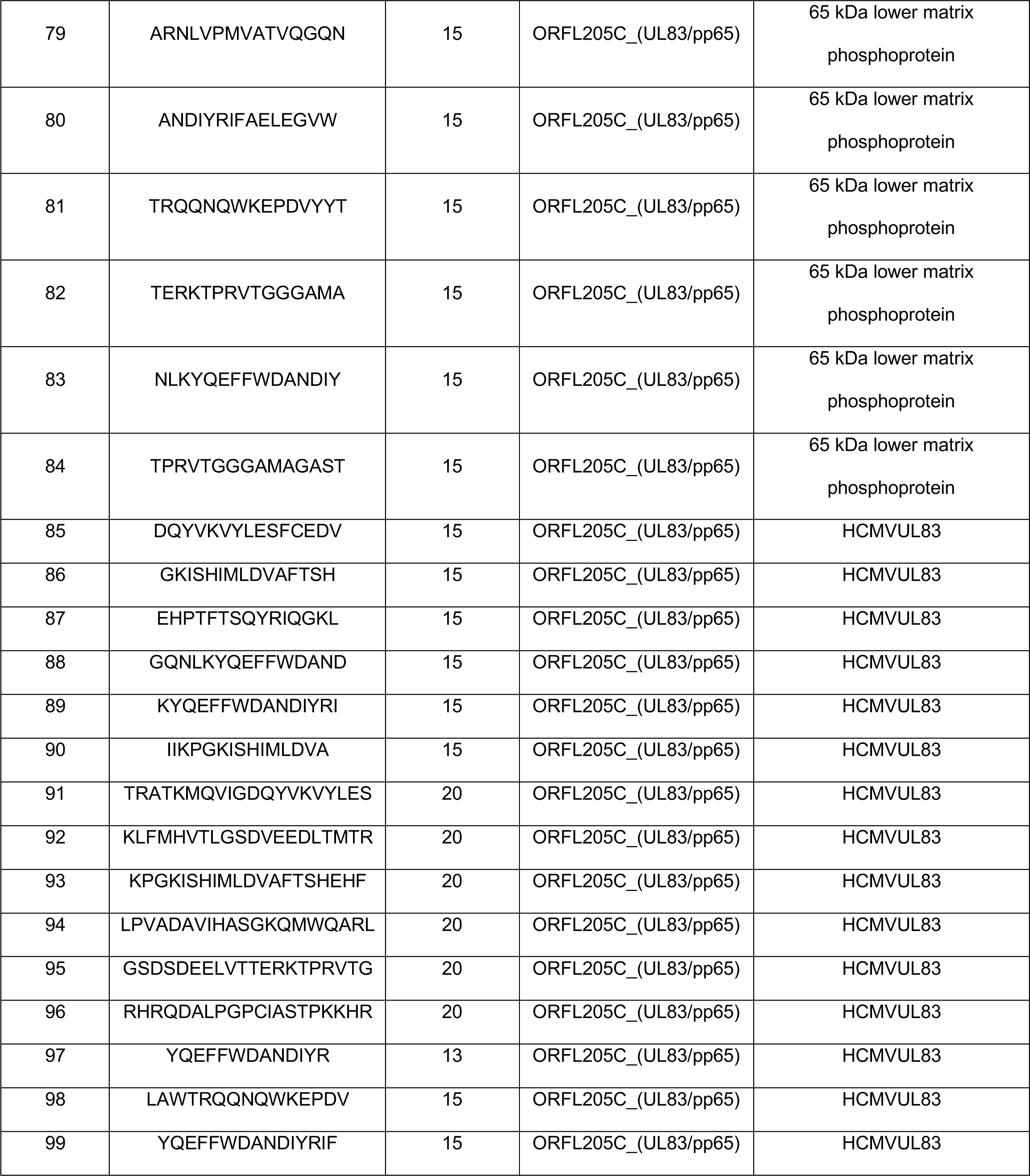
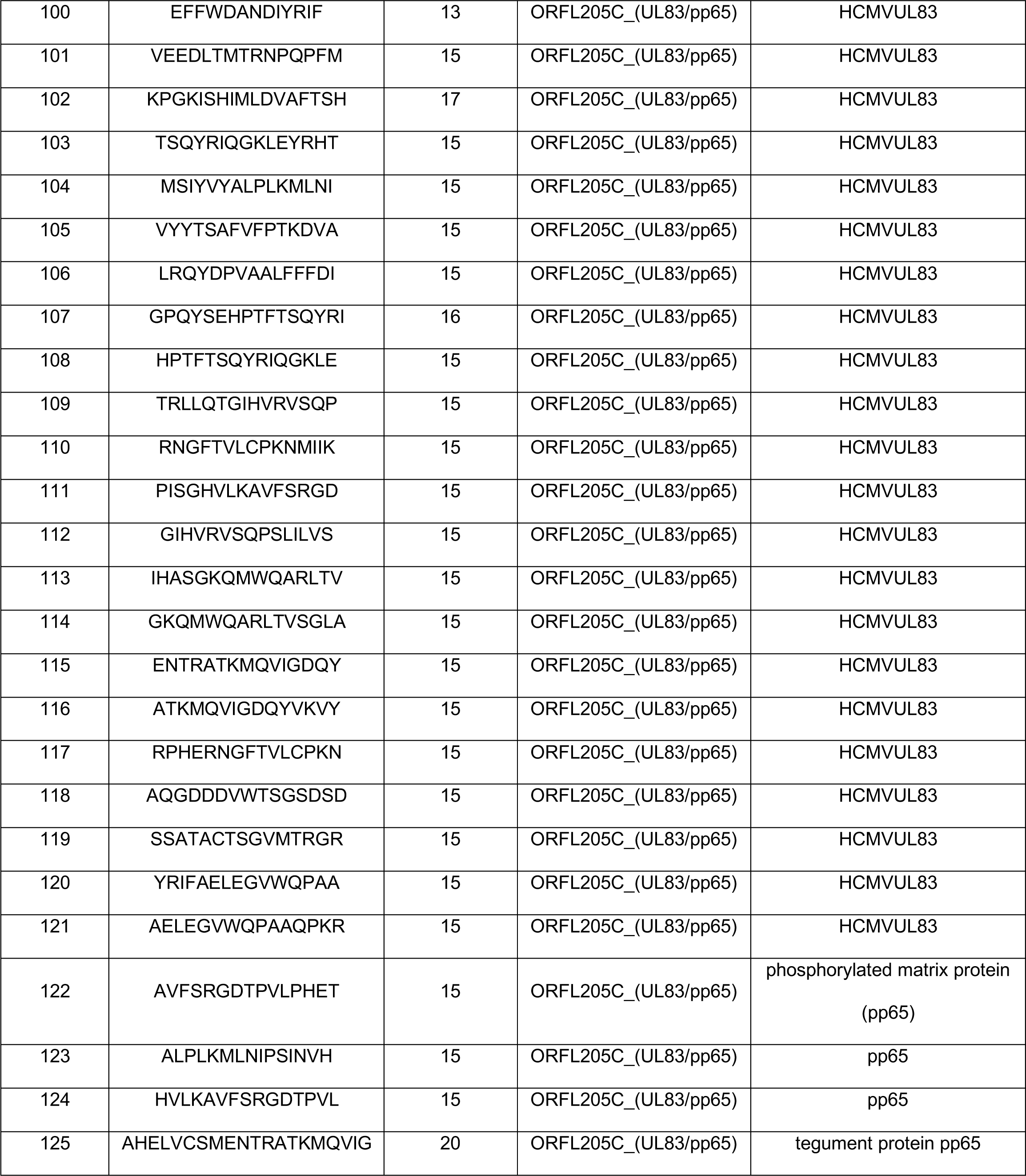
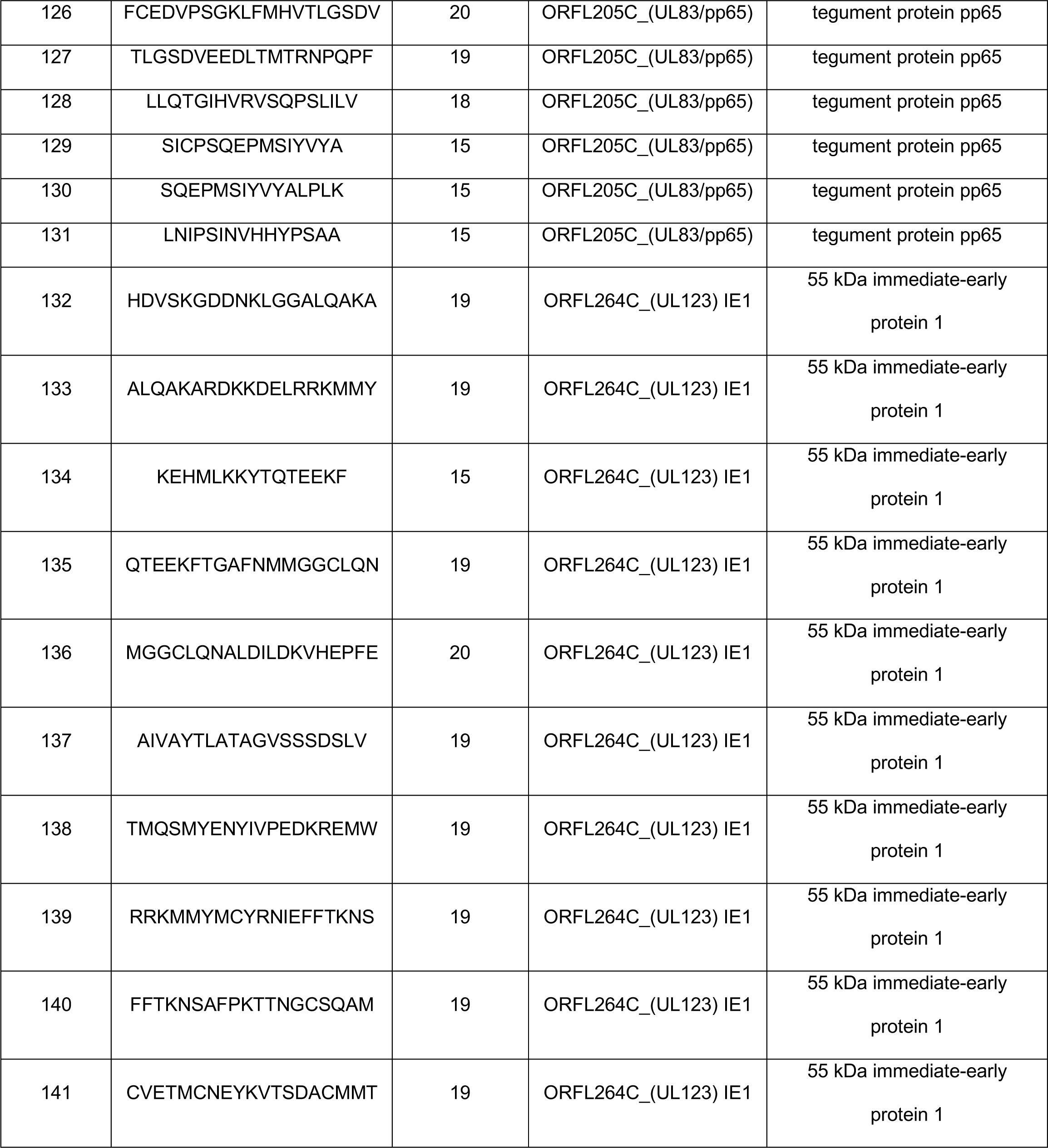
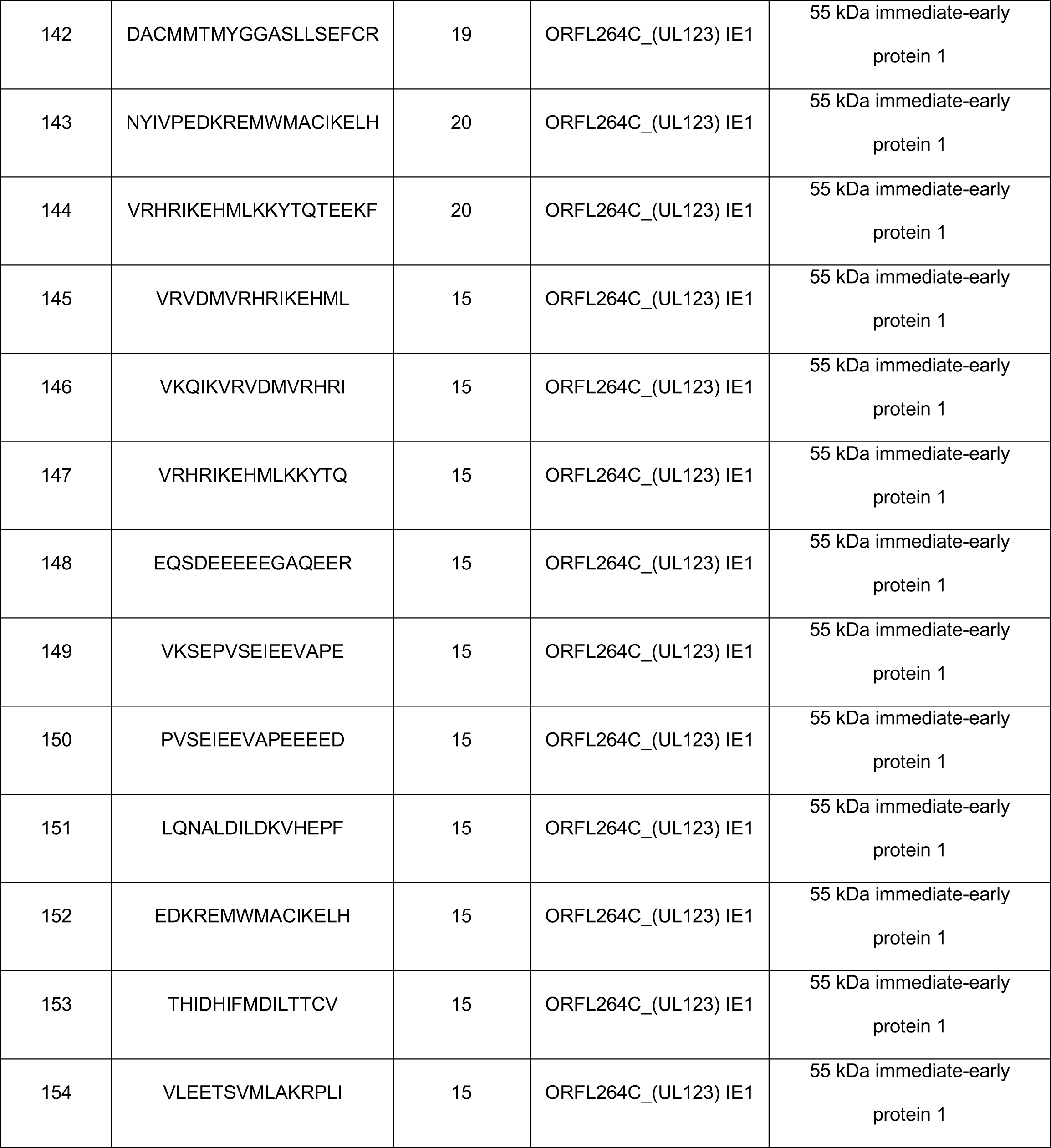
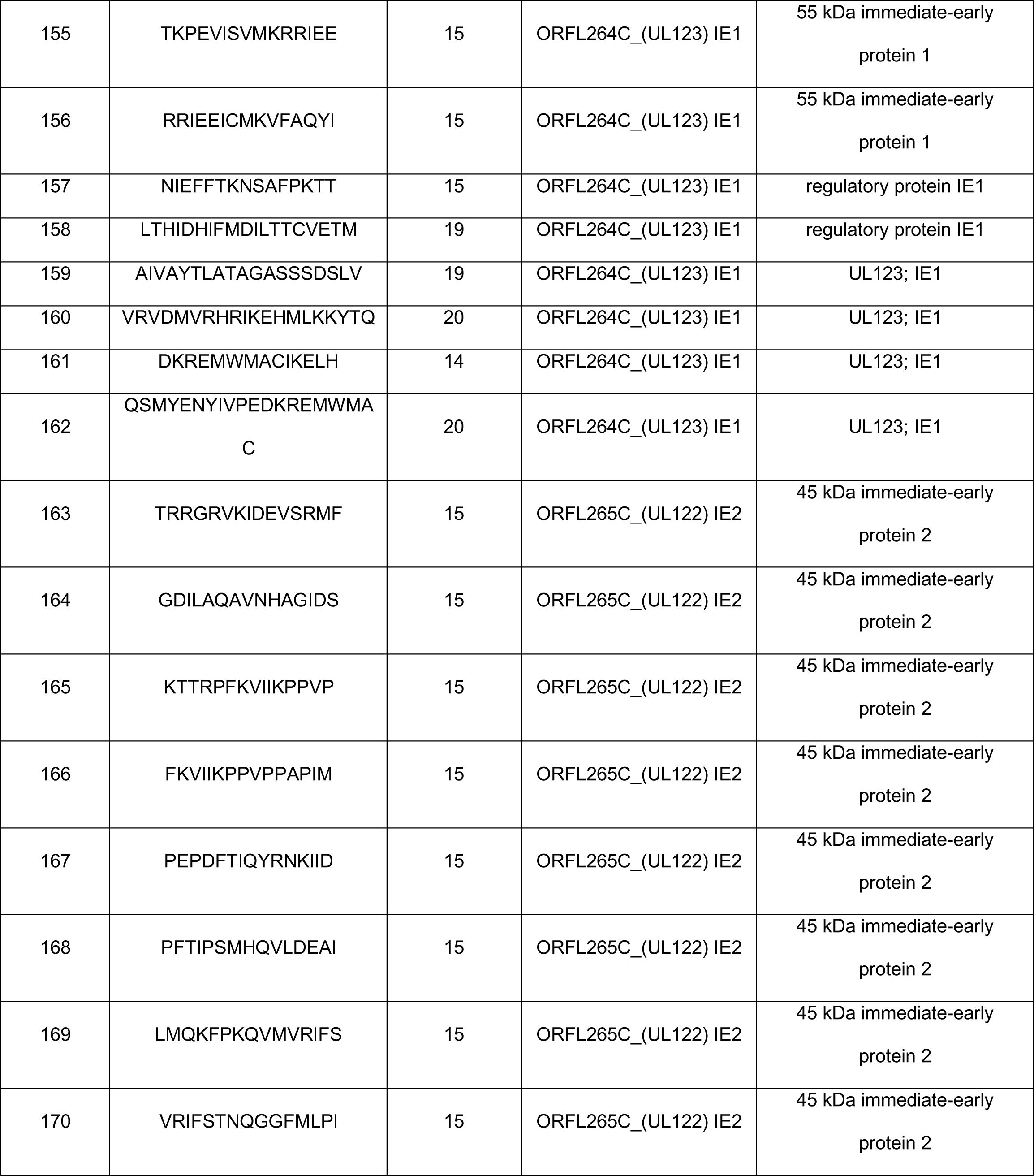
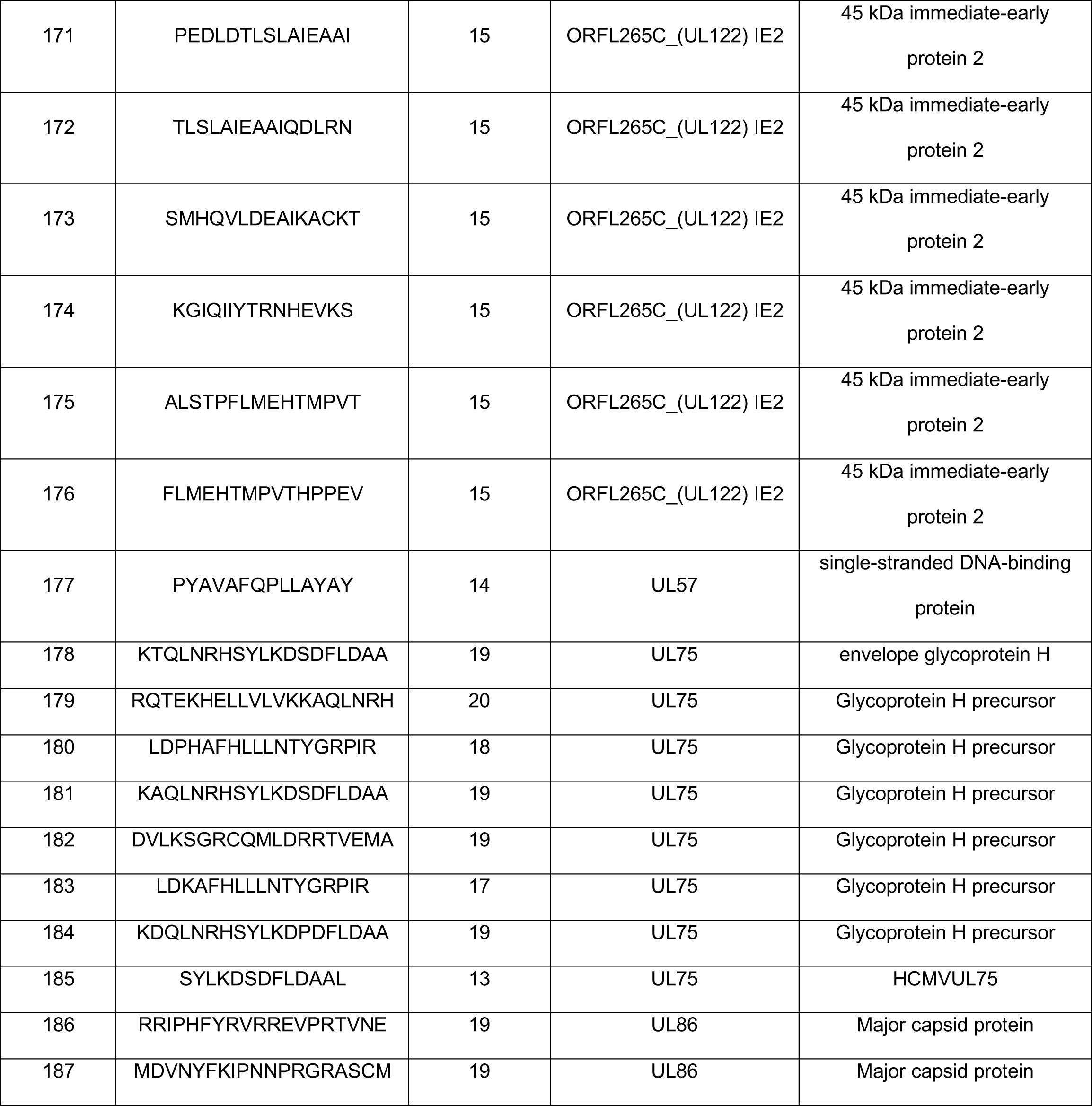
Details of HCMV specific class II epitopes from IEDB

These MPs were tested with PBMC from a new cohort of 20 individuals (6 males and 14 females), which included both HCMV seropositive and seronegative donors (10 CMV^+^ and 10 CMV^-^, **Fig. S1B** for IgG ELISA CMV confirmation). None of the PBMC from these subjects were used in the original epitope mapping experiments. PBMCs were stimulated with the Mabtech, P235, IEDB-II, or a combination of both P235/ IEDB-II MPs. CD4+ T cell responses were measured as percentage of activation-induced marker assay positive (OX40+ CD137+) CD4+ T cells and results are displayed in **Fig. 6** (flow cytometry gating strategy shown in **Fig. S5B**).

**Fig. 6.**
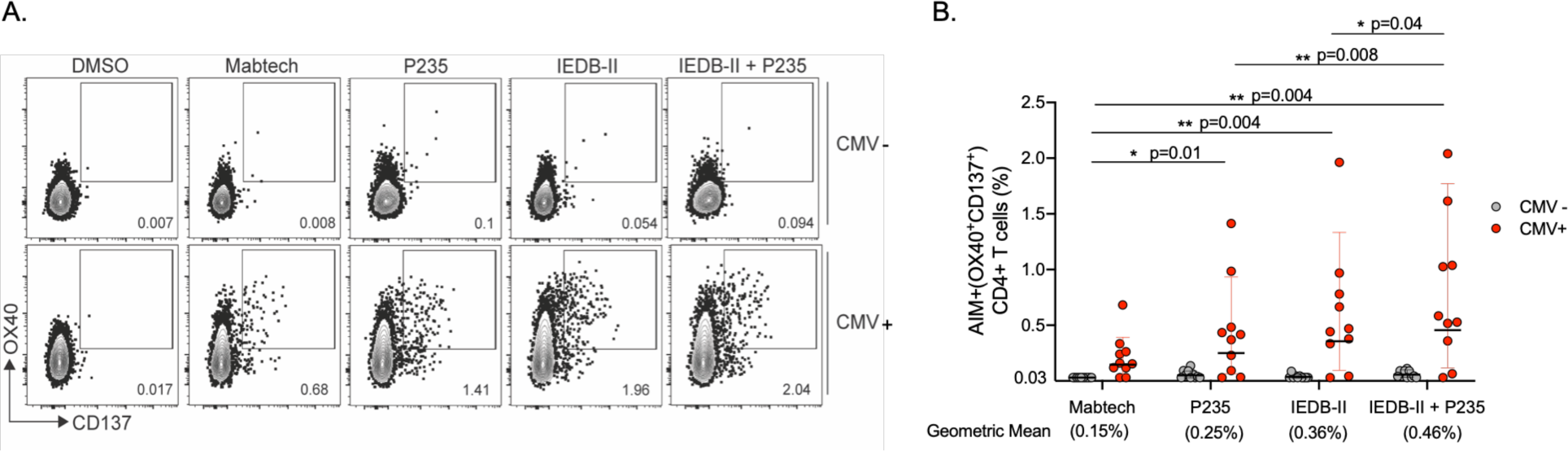
Epitope specific CD4+ T cell responses in HCMV+ and HCMV-subjects detected with different peptide pools: (A) Representative FACS plots showing HCMV specific CD4+ T cell reactivity against different peptide pools based on activation-induced marker assays (OX40+ and CD137+ double expression). PBMCs from HCMV+ (red circles) and CMV-donors (grey circles) were stimulated with 2 µg/ml of the Mabtech pool or IEDB-II/P235 pools for 24 hrs. (B) Epitope-pool specific CD4+ T cells measured as percentage of activation-induced marker assay positive (OX40+ CD137+) CD4+ T cells. Each dot represents an individual subject. HCMV+ subjects demonstrated significantly higher CD4+ T cell AIM responses than HCMV-subjects with all the different pools tested. Mabtech HCMV+ vs HCMV- p=0.0007; P235 HCMV+ vs HCMV- p=0.0065; IEDB-II CMV+ vs CMV- p=0.0009; P235/IEDB-II CMV+ vs CMV- p=0.004. Two-tailed Mann-Whitney test. Comparisons across different pool formulations within the CMV+ were made using the Wilcoxon matched-pairs signed ranked test, Two-tailed p values are shown in the Figure; Geometric mean with geometric standard deviation.

All HCMV MPs tested were associated with significantly higher CD4 AIM responses in HCMV+ individuals compared to HCMV-subjects as shown in **Fig. 5** (statistical differences detailed in figure legend). When comparing AIM responses between the HCMV pools, the P235, IEDB-II and P235/IEDB-II MPs were associated with significantly higher HCMV-specific CD4 responses compared to the Mabtech pool (geometric mean 0.15% vs 0.25% CD4 AIM+, p=0.01; and 0.15% vs 0.36%, p=0.004, and 0.15% vs 0.46% CD4 AIM+, p=0.004, respectively by Wilcoxon test). This was expected, as the Mabtech pool contains fewer epitopes which are also mainly CD8 T cell specific. Additionally, the combination of the P235 and IEDB-II MPs elicited higher CD4 responses than either MP alone (geometric mean 0.25% vs 0.46% CD4 AIM+, p=0.0078 and 0.36% vs 0.46% CD4 AIM+, p=0.004, respectively by Wilcoxon test) and had the highest magnitude response of all pools tested. This indicates that the combination of known (IEDB-II MP) and novel epitopes and ORFs (P235 MP) can capture the broadest range of CD4 T-cell responses in HCMV+ individuals, which has high potential for clinical diagnostic use.

## Discussion

In this study we have identified >200 new epitopes derived from 100 HCMV ORFs that induce virus-specific T cell responses. Importantly, this demonstrates that the current HLA peptide-binding prediction algorithms that we and others have refined over the last several decades are extremely efficient (47–51), and represent an excellent alternative to synthesizing genome-wide overlapping peptides, especially for large pathogens such as HCMV. Despite the significant diversity in the human HLA repertoire, current advances in algorithm-based epitope identification take into consideration epitopes with potential binding to diverse haplotypes, which undoubtedly contributed to this success (40, 52). Together, this approach allowed us to increase the known T cell epitope landscape for HCMV by greater than 10-fold by synthesizing only 2593 peptides, illustrating both its efficiency and cost effectiveness in deciphering immune targets of large pathogens.

We chose to use IFN-γ production as a readout for positive epitope reactivity in a fluorospot-based assay to identify HCMV-specific T cell epitopes in this study. As true for most viral infections, CMV drives a strong Th1-like CD4+ response, and most effector and memory viral CD8+ T cells also produce this cytokine (53). However, future studies assessing which of these 235 epitopes may elicit HCMV-specific CD4 T cells to produce other cytokines are merited. Previously, we have observed that Dengue virus epitope-specific CD4+ T cells can produce both IFNγ and IL-10 (54), something we have also seen during acute CMV infection in mice (55), where IL-10 producing CD4+ T cells enhance the duration of viral persistence (56). Recent studies by the Wills and Moss groups show that subsets of HCMV epitope-specific CD4+ T cells can produce IL-10 and also display cytolytic markers (57, 58). The potential CTL activity of HCMV-specific CD4+ T cells has been postulated for many years (59), and our recent results showing that CMV epitope-specific CD4 T cells can directly kill *in vivo* support this hypothesis (60). Taken together, our identification of >200 new T cell epitopes that elicit IFNγ production in this study provide us and others in the field valuable new tools to dissect the phenotypes and effector functions of HCMV-specific CD4 T cells in cases of both healthy and immune compromised patients, and will also help instruct ongoing vaccine efforts.

Of the 100 ORFs which we show here to be sources of specific T cell epitopes, 41 were uniquely identified as ribosome-bound RNAs in HCMV infected fibroblasts (39), with these 41 yielding 50 unique epitopes. Notably, of these 41 ORFs, 17 are predicted to produce proteins <50 amino acids in length, and 7 contain non-ATG start codons. This is consistent with recent studies suggesting that the short/‘cryptic’ mRNAs present in both virally infected and tumor cells can be translated, proteolytically processed and loaded onto HLA molecules, resulting in the induction of epitope-specific T cell responses (61–63). Interestingly, one of the larger 41 ORFs that contains two newly identified T cell epitopes (ORFL147C, 476 amino acids) has very recently been shown to regulate RNA binding/processing, and its deletion compromises CMV replication in fibroblasts (64). Despite >20% of the novel T cell epitopes identified here being derived from these newly described, ribosome-associated HCMV RNAs, no more than 2 of the 19 healthy donors analyzed produce T cells specific for any single one of these epitopes. This indicates that these novel ORFs 1) may not be broad targets of T cell responses in infected persons, 2) that specific individuals may more efficiently present epitopes derived from short/cryptic HCMV RNAs or 3) that minor HLA molecules may present them, with other possibilities also existing. Additionally, whether the proteins derived from these short ORFs are stable and play a role in the HCMV lifecycle remains an open question. Finally, we also identified 24 epitopes derived from 14 ‘canonical’ HCMV ORFs where the only historic support for their existence was the presence of their RNA in infected cells or bioinformatic analyses. Notably, a recent comprehensive study where 169 predicted canonical HCMV proteins (including these 14) were epitope-tagged, expressed stably in infected cells, immunoprecipitated and analyzed for interacting proteins by mass spectrometry supports our results that these ORFs are expressed as proteins (64).

Of the 59 canonical ORFs that we have identified here to contain T cell epitopes, >25% of these are known to function as immunomodulatory proteins (65). This is intriguing, as perhaps these HCMV proteins are more subject to being localized to antigen-processing or presentation compartments within infected cells. One of these epitopes is derived from the HCMV IL-10 orthologue, which is being considered as a potential HCMV vaccine candidate (66, 67). Additionally, 3 epitopes were found to be embedded within the viral UL128 protein, a critical component of the pentameric envelope protein complex (UL128-131/gH/gL) that mediates entry of HCMV into non-fibroblast cell types (68, 69). This is also of high potential interest in the context of vaccine development, as many believe the pentamer should be included in a viral- or subunit-based approach (70). Notably, both vIL-10 and UL128 have largely been considered only in the context of their abilities to induce antibody-based vaccine protection, but our identification of T cell epitopes derived from both these HCMV proteins suggests they may function to prime both humoral and cellular immunity.

## Methods

### Study design

For the initial CMV ORF screen, the responses of 19 CMV-seropositive subjects were evaluated. PBMCs were stimulated with 89 pools covering 563 ORFs of HCMV. Each pool comprised of 28-30 15-mer peptides overlapping by 10 residues. PBMCs that were found reactive to a pool were further tested against individual peptides contained in the pool using IFN-γ Fluorospot assay. Flow cytometry was then used to further characterize the epitopes recognized by PBMCs stimulated with individual peptides by detecting IFN-γ production from CD8+ and CD4+ T cells.

For the CMV-235 validation and comparison screen, the responses of a new cohort consisting of 10 CMV-seropositive and 10 seronegative subjects were evaluated. PBMCs were stimulated with CMV-Mabtech peptide pool (Catalog 3619-1), CMV-IEDB peptide pool (Table 2) (44, 46), CMV-235 pool, or a combination of both CMV-IEDB and CMV-235 pools. PBMC responses were assayed using the same IFN-γ Fluorospot assay. These studies were approved by the institutional review board committee at La Jolla Institute protocol number: VD-112 and VD-174.

### Subjects

19 subjects (10 males and 9 females) were recruited anonymously from San Diego blood bank (SDBB) for the initial CMV ORF screens. For the CMV-235 comparison screens, samples from 20 subjects (6 males and 14 females) were obtained by La Jolla Institute Clinical Core and Continental Services Group (Miami, FL) for prior, unrelated studies. Blood samples were collected by trained staff. At the time of enrollment in the initial studies, all individual subjects provided informed consent that any leftover sample could be used for future studies, which includes this study. These subjects were considered healthy as defined by no known history of any significant systemic diseases (not limited to autoimmune disease, diabetes, kidney or liver disease, congestive heart failure, malignancy, coagulopathy, hepatitis B or C, or HIV). The demographics of those subjects are provided in **Table 3**.

**Table 3:** Demographic characteristics of HCMV (+/-) subjects analyzed in screening and validation studies.

| **Characteristics** | *Screening cohort* | *Validation cohort* |  |
| --- | --- | --- | --- |
|  | **HCMV+** | **HCMV+** | **HCMV-** |
| Total participants enrolled, n | 19 | 10 | 10 |
| Males/females | 10/9 | 3/7 | 3/7 |
| Median age (range) | 65 (28-80) | 35.5 (22-55) | 28.5 (19-42) |
| Caucasian, % (n) | 68 (13) | 40 (4) | 40 (4) |
| Asian, % (n) | 16 (3) | 10 (1) | 20 (2) |
| African American, % (n) | 5 (1) | 10 (1) | 10 (1) |
| More than one race, % (n) | 0 (0) | 30 (3) | 30 (3) |
| Unknown, % (n) | 10 (2) | 10 (1) | 0 (0) |

The IgG antibodies of the subjects for both cohorts were measured using Cytomegalovirus IgG Elisa kit from Genway Biotech Inc. according to manufacturer’s instructions.

### Peptide prediction

Based on the 7-allele method as previously described (40), 2593 peptides were predicted for 563 potential HCMV ORFs. Of the 751 ORFs predicted by ribosomal profiling (39), those smaller than 15 amino acids were excluded, and only one peptide of ORFs 15-20 amino acids in length were selected for screening.

### Peptide libraries and pool preparation

The predicted peptides were commercially synthesized as crude material by TC Peptide Lab (www.tcpeptidelab.com; San Diego, CA). The peptides were solubilized in DMSO at a concentration of 20 mg/ml and spot checked for quality by mass spectrometry. The peptides were pooled into peptide pools containing 28-30 peptides constituting multiple ORFs per pool. A total of 89 pools were prepared covering 563 ORFs of HCMV. The final concentration of each pool was 0.7 mg/ml.

For the IEDB-II (Table 2) and P235 (Table 1) peptide pools peptides were synthesized by A&A ltd, San Diego, resuspended in DMSO, pooled and sequentially lyophilized as previously described (71). The IEDB-II peptide pool was developed based on data available in the IEDB (www.iedb.org) (41). The MHC class II restricted epitopes for CMV was extracted from the IEDB in October of 2020 using the following query; Organism: human herpesvirus 5 (ID:10359), positive assays only, no B cell assays, MHC restriction type: class II, host: *Homo sapiens*. The resulting 187 epitopes (table 2) were filtered for size (13-20 amino acids) and discovered using one of the following assays: ELISPOT, ICS, multi- or tetramers, proliferation and “helper response”. The CMV peptide pool for human CD4 and CD8 T cells containing 42 peptides (14 MHC class II restricted and 28 MHC class I restricted) representing pp50, pp65, IE1, IE2, and envelope glycoprotein B was purchased from Mabtech.

### Isolation of PBMC by Ficoll-Paque density gradient centrifugation

One-unit blood from each donor was processed for PBMC isolation. Briefly, blood was centrifuged and the top layer of plasma was removed. The remaining blood was diluted and layered over 15 ml of Ficoll-Paque. Tubes were spun at room temperature in a swinging bucket rotor without brake applied. The PBMC interface was carefully removed by pipetting and washed with PBS by centrifugation at 800 rpm for 10 mins with brakes off. PBMC pellet was resuspended in RPMI media, cell number and viability were determined by trypan blue staining and cells were cryopreserved in liquid nitrogen in freezing media (90% Fetal bovine serum and 10% DMSO) at a density of 30 million/ml and stored until further processed.

### Fluorospot assay

PBMC were thawed, washed and counted for viability using the trypan blue exclusion method. 200,000 cells were plated in triplicates and stimulated with pools (2µg/ml) or peptides (10µg/ml), PHA (10µg/ml) or medium containing equivalent amount of DMSO in 96- well plates (Immubilion-P, Millipore) previously coated with anti IFN-γ antibody (1-D1K, Mabtech, Stockholm, Sweden). After 20 hr incubation at 37°C, cells were discarded and wells were washed six times with PBS/0.05% Tween 20 using an automated plate washer and further incubated with IFN-γ antibody (7-B6-1-FS-BAM) for 2 hrs at room temperature. After incubation, wells were washed and incubated with fluorophore conjugated anti-BAM-490 antibody for 1 hr at room temperature. Finally, the plates were washed and incubated with fluorescence enhancer for 15 min, blotted dry and fluorescent spots were counted by computer assisted image analysis (IRIS Fluorospot reader, Mabtech, Sweden).

Each pool or peptide was considered positive compared to the background that had equivalent amount of DMSO based on the following criteria: (i) 20 or more spot forming cells (SFC) per 10^6^ PBMC after background subtraction, (ii) the stimulation index greater than 2, and (iii) p<0.05 by student’s t test or Poisson distribution test when comparing the peptide or pool triplicates with the negative control triplicate.

### Intracellular cytokine assay for IFN-γ

Intracellular staining for IFN-γ and flow cytometry was performed to detect antigen specific T cell responses. 1x10^6^ PBMCs suspended in RPMI medium supplemented with 1-% heat inactivated human AB serum, glutamine and penicillin streptomycin were plated in U-bottom 96 well plates. After overnight resting at 37°C, PBMCs were spun and replaced with fresh RPMI media and stimulated with individual peptides at a concentration of 10 µg/ml. PHA at a concentration of 5 µg/ml was used as a positive control. After 1 hr of incubation at 37°C, 2μg/ml of Brefeldin was added and cell were further incubated at 37°C for additional 5 hrs. The cells were then harvested, washed with 200 µl of MACS Buffer and stained with a cocktail of antibodies that contained CD3-Af700 (eBioscience, clone UCHT1), CD4-APCef780 (eBioscience, clone RPA-T4), CD8-BV650 (Biolegend, clone RPA-T8), CD14-V500 (BD Biosciences, clone M5E2), CD19-V500 (BD Biosciences, clone HIB19), and fixable viability dye-e506 for 30 min at 4°C. The cells were then washed thrice with 200 µl MACS buffer, fixed using 4% PFA for 10 mins at 4°C, washed with 200 µl PBS and rested at 4°C overnight in 200 µl MACS buffer. The following day, cells were washed, permeabilized by washing with 200 µl saponin buffer (0.5 % saponin in PBS), washed with blocking buffer (10% human serum prepared in saponin buffer) and stained with IFN-γ-FITC (eBioscience, clone 4S.B3) antibody at room temperature for 30 mins. The cells were finally washed with PBS and suspended in 200 µl PBS.

The cells were acquired on ZE5 Biorad plate reader and further analysis was done on FlowJo software. Gates were applied on live single cells for CD3+, CD4+ and CD8+ T cell populations. The percentage of reactive CD4+ or CD8+ IFN-γ T cells were expressed as a percent of the total number of parent population analyzed. Reactive populations met the following 2 criteria: (i) well-defined cell population positive for both IFN-γ and CD4 or CD8 constituting at least 0.02% (post subtracting their corresponding DMSO controls) of the total number of CD4+ or CD8+ cells analyzed (ii) stimulation index greater than 2.

### Activation induced marker (AIM) assay

PBMC were thawed, washed and counted for viability using the trypan blue exclusion method. 1 million cells per donor/condition were plated and cultured in the presence of the CMV specific pools (1µg/mL for P235 and IEDB-II, 2µg/mL for Mabtech pool), PHA (10µg/mL), or medium containing equivalent amount of DMSO in 96-well U-bottom plates. Cells were then harvested, washed with 200µl of MACS Buffer and stained with a cocktail of antibodies that contained CD3-Af700 (eBioscience, clone UCHT1), CD4-BV605 (eBioscience, clone RPA-T4), CD8-PerCP-Cy5.5 (Biolegend, clone HIT8a), CD14-V500 (BD Biosciences, clone M5E2), CD19-V500 (BD Biosciences, clone HIB19), OX40-PE-Cy7 (Ber-ACT35), CD137-APC (4B4-1), and fixable viability dye-e506 for 30 min at 4°C. The cells were then washed thrice with 200 µl MACS buffer, fixed using 4% PFA for 10 mins at 4°C, and resuspended in 200 µl of PBS for acquisition.

Cells were acquired on a BD LSRFortessa and further analysis was done on FlowJo software. As previously described (44, 72), quantification of live, singlet antigen specific CD4 T cells was determined as a percentage of their OX40+CD137+ expression (AIM+). CMV specific AIM+ CD4 T cell signals were background subtracted with their corresponding negative control DMSO samples, with a minimal DMSO level set to 0.005%. The limit of detection (LOD) for the AIM+ assay was calculated by multiplying the upper confidence interval of the geometric mean of all DMSO samples by 2 (0.03).

### Statistical analysis

Statistical analyses were performed using GraphPad Prism versions 8.1.1 and 8.4.3. Statistical details are provided with each figure.

## Acknowledgement

We would also like to thank all donors that participated in the study. We also thank the La Jolla Institute for Immunology Clinical Studies Group and Flow Cytometry Core for all the invaluable help.

## Author contribution

AS and CAB conceived the study. RD and GW performed the experiments and analyzed the data. SKD performed peptide prediction, JP processed blood samples, RD, GW, and GP conducted ELISA, JS helped with the quality check of synthesized peptides, AG designed the IEDB-II pool, CLA, AS, CAB directed the study, RD, AS, CAB wrote the manuscript taking input from other authors.

## Supplementary figures

**Fig. S1.**
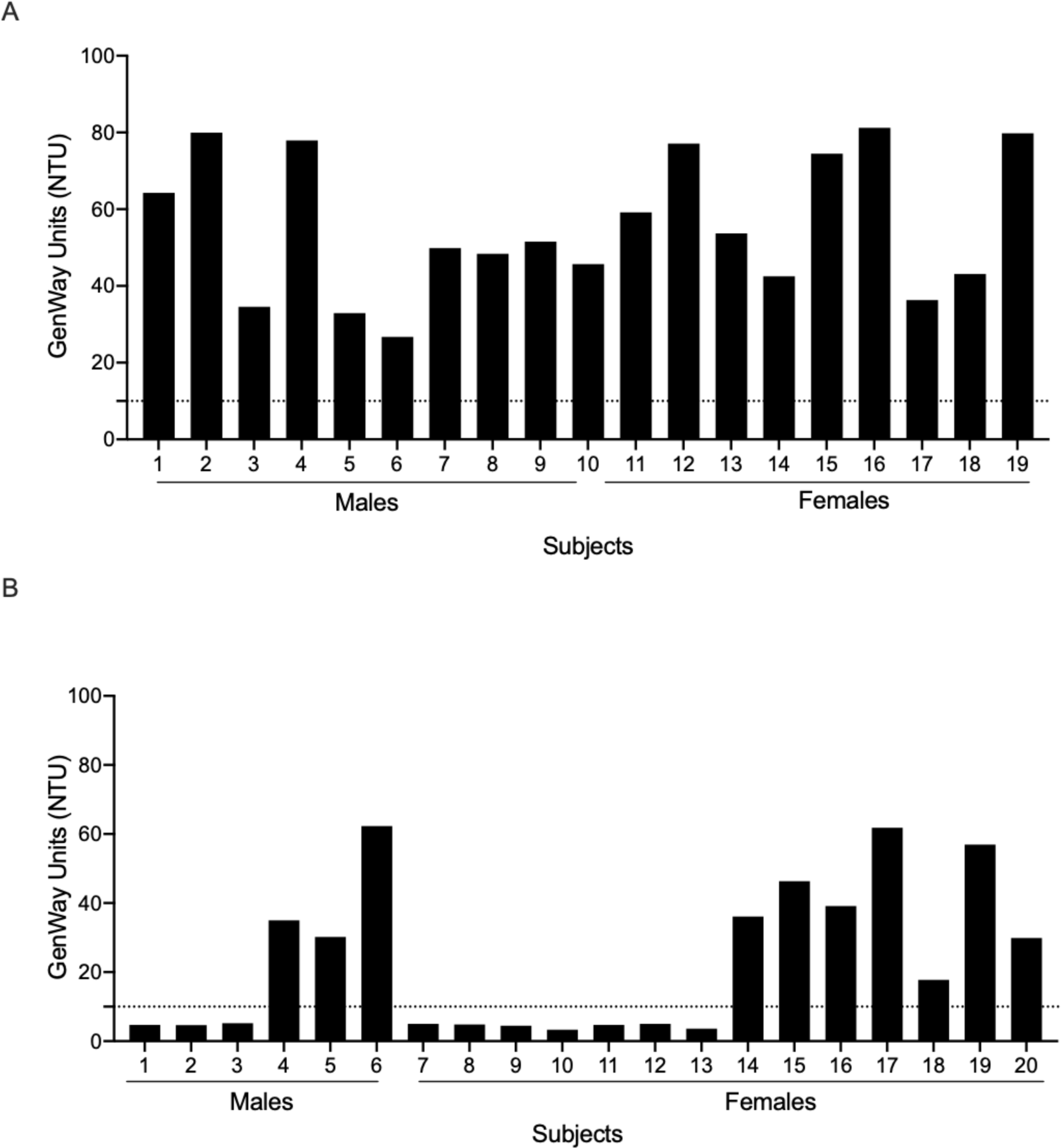
Confirmation of HCMV seropositivity in donors. (A) IgG levels in plasma of subjects in the screening cohort (n=10 males, n=9 females) and (B) the validation cohort (n=13 males, n=26 females) determined by ELISA. Dotted line represents the cut off for positivity (10 NTU).

**Fig. S2.**
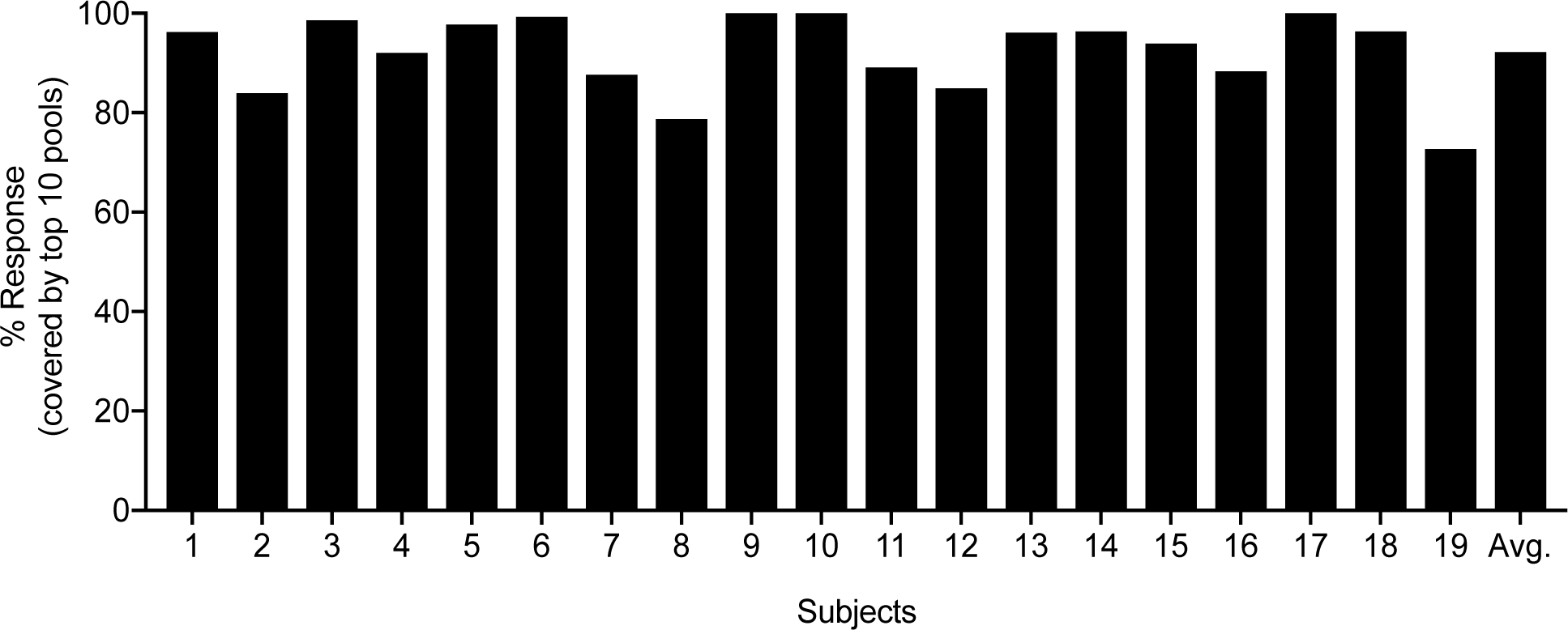
The total T cell response captured by the top 10 epitope pools in each subject. The response magnitude of the top 10 pools as a percentage of the total response magnitude observed from all positive pools. On average, the top 10 pools accounted for ∼90 % of each subject’s total response.

**Fig. S3.**
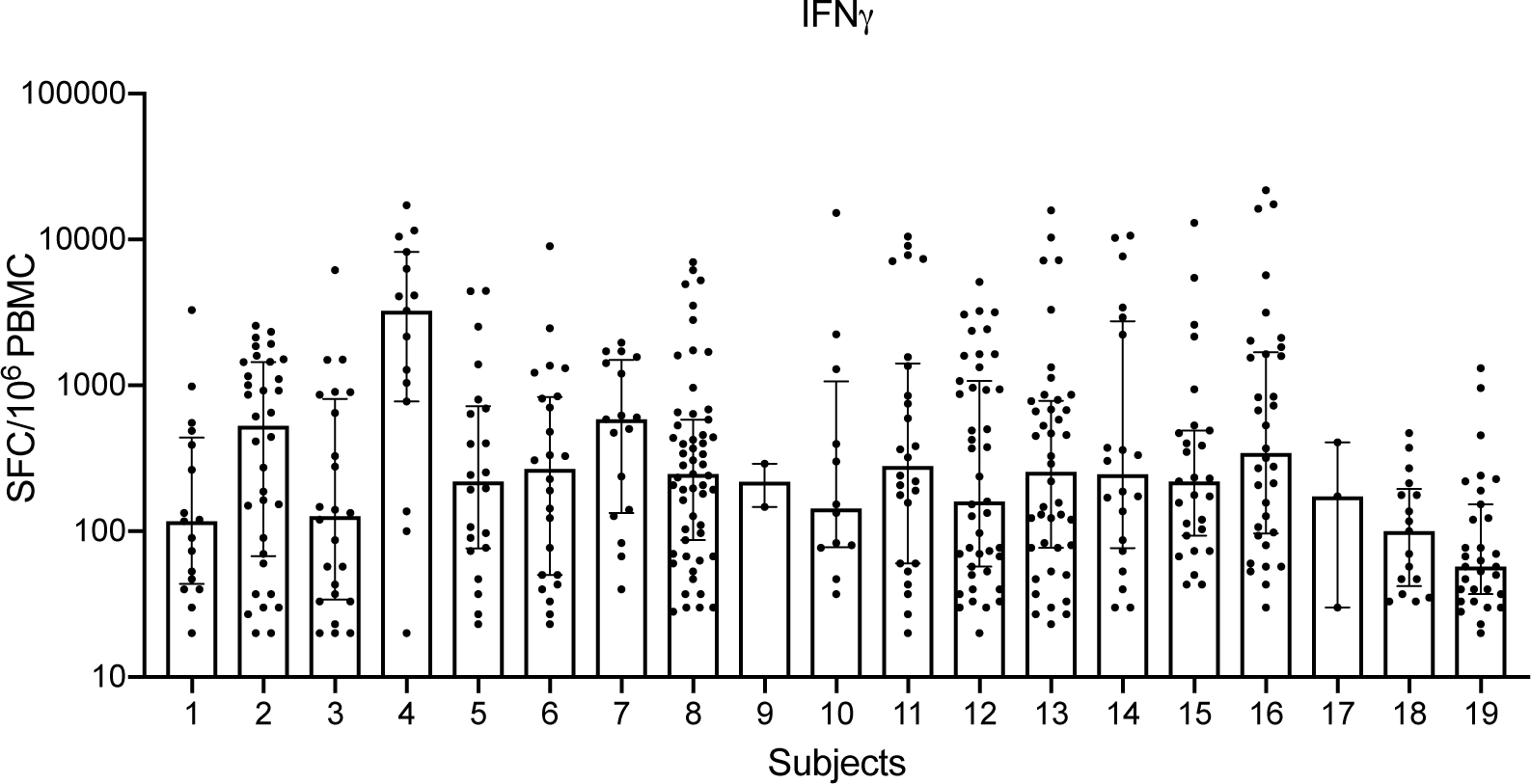
Response magnitude of each epitope identified in HCMV seropositive individuals: Each dot represents an epitope. Y axis represents the response magnitude of individual epitopes. X axis represents each subject. Median ± interquartile range is shown.

**Fig. S4.**
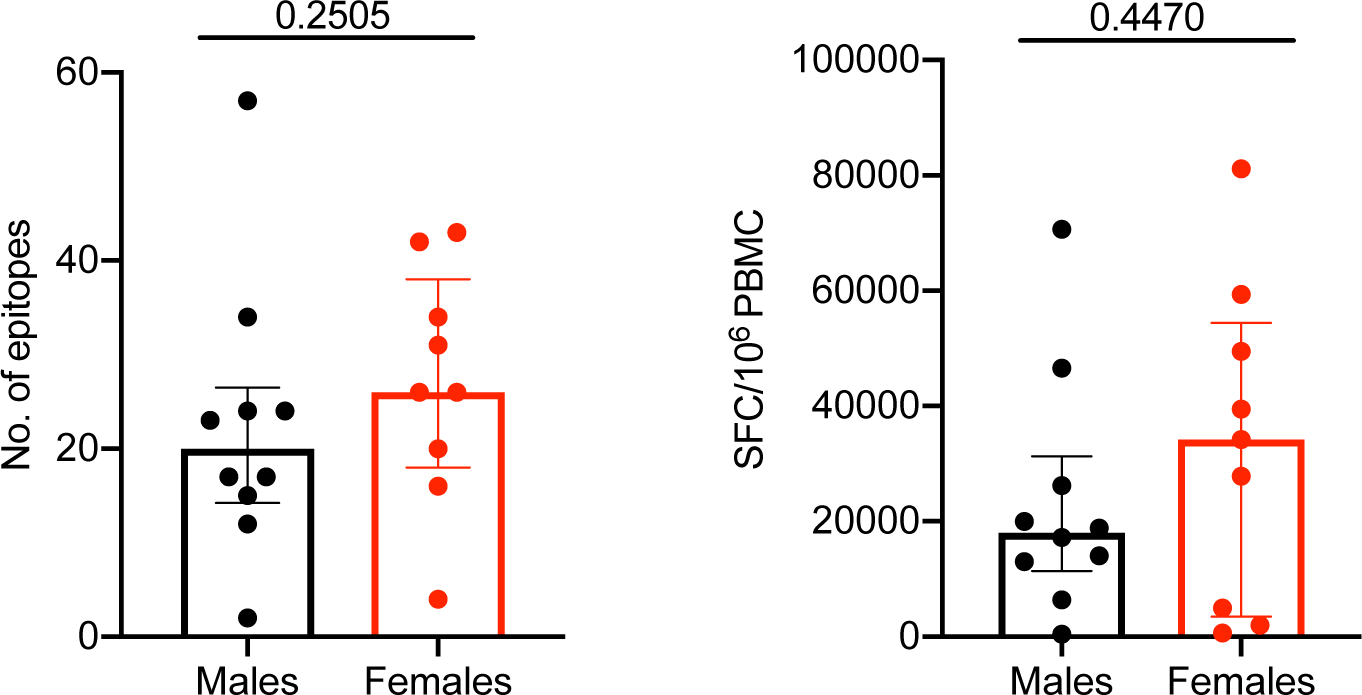
Frequency and magnitude of response in males and females: Each dot represents a donor. Black dot/bar represents males and red dot/bar represents females. Median with interquartile range is displayed. Two-tailed Mann-Whitney test.

**Fig. S5.**
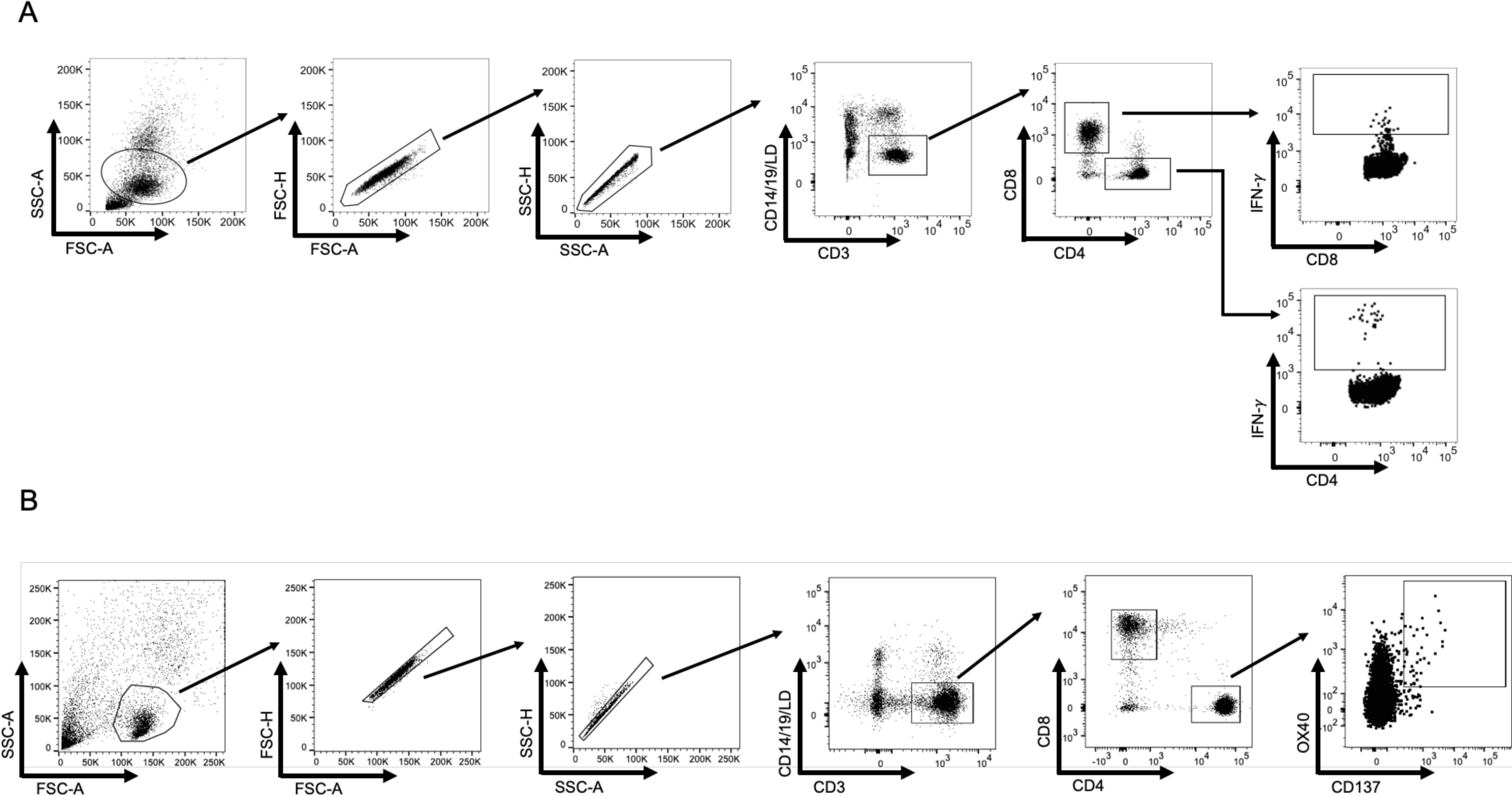
Gating strategy adopted in IFN-γ Fluorospot and AIM assay: (A) Human PBMCs isolated from HCMV+ subjects were stimulated with each scoring peptide to identify HCMV-specific IFN-γ producing CD4+ and CD8+ T cells. (B) Human PBMCs isolated from HCMV+ and HCMV-subjects were stimulated with each megapool generated to identify HCMV-specific activation-induced marker assay positive (OX40+ CD137+) CD4+ T cells.

